# SARS-CoV-2 Main Protease Inhibition Through Dimerization Promotion by Peptidomimetic Inhibitors and Disruption by Ebselen

**DOI:** 10.1101/2025.11.16.688752

**Authors:** Chengxi Liu, Qinyu Jia, Chang Zhao, Zhong-Ping Yao

## Abstract

The SARS-CoV-2 main protease (M^pro^) is a key antiviral target that equilibrates between an active dimer and inactive monomer, making the dimerization interface a promising therapeutic vulnerability. This study investigates M^pro^ monomer-dimer equilibrium and conformational changes induced by inhibitor binding using ^13^C labeling with native mass spectrometry and hydrogen/deuterium exchange mass spectrometry (HDX-MS). We evaluated how peptidomimetic inhibitors (PF-07321332, PF-00835231, GC376, boceprevir) and non-peptidomimetic inhibitors (carmofur, ebselen, MR6-31-2, AT7519, pelitinib) influence M^pro^ dimerization and subunit exchange. Key findings revealed divergent mechanisms: peptidomimetic inhibitors significantly shifted the equilibrium towards the dimeric state, suppressing subunit exchange dynamics and rigidifying the dimer interface; in contrast, ebselen impaired dimer form and increased interface flexibility. Notably, tandem mass spectrometry identified a novel covalent binding site for ebselen at C300. Molecular dynamics simulations and mutational analyses demonstrated that this C300 modification allosterically alters the hydrogen bond network of the dimer interface, directly contributing to ebselen-mediated dimer disruption and enzymatic inhibition. Overall, this study clarifies distinct inhibitory modes between peptidomimetics and ebselen, highlighting the therapeutic potential of targeting allosteric sites at the dimer interface for designing next-generation M^pro^ inhibitors.

## Introduction

Severe acute respiratory syndrome coronavirus 2 (SARS-CoV-2), the causative agent of COVID-19, caused global infections, fatalities, and profound socioeconomic disruptions (Zhu et al., 2020). SARS-CoV-2 contains a ∼30 kb single-stranded RNA, which includes the ORF1ab gene that encodes two large overlapping polyprotein precursors (pp1a and pp1ab) (Zhang et al., 2020). The cleavage of the two inactive polyproteins into 16 individual active nonstructural proteins is essential for viral replication and proliferation. This cleavage process is performed by two viral proteases: the papain-like protease and the main protease (M^pro^). M^pro^ is a cysteine protease produced by autolytic cleavage from pp1a and pp1ab and is responsible for the generation of 12 nonstructural proteins from the two polyproteins (Yang et al., 2003). M^pro^ exhibits a highly conserved three-dimensional structure among various coronaviruses, e.g., SARS-CoV-2, SARS-CoV, MERS-CoV, and BAT-CoV (Goyal and Goyal, 2020), offering the potential for broad-spectrum anti-coronaviral drug development.

M^pro^ is a homodimeric protein, with each monomer consisting of three domains. Domain І (residues 10-99) and Domain II (residues 100-182) feature an antiparallel β-barrel structure, and Domain III (residues 198-303) contains five α-helices connected to Domain II by a long loop linker (residues 183-197) (Jin et al., 2020a). The substrate-binding site (active site), located in a cleft between Domain I and Domain II, has been defined into four subsites (S1, S2, S4 and S1’), which contain a histidine/cysteine catalytic dyad: H41 and C145 (Figure 1a). SARS-CoV-2 M^pro^ is in an equilibrium between the active dimer and inactive monomer in solution. Dimerization is essential for the M^pro^ to perform enzymatic activity, with key dimer interface residues playing a critical role in shaping the structure of the active site pocket. The dimer interface includes the N-finger (residues 1-7), residues 121-129, residues 132-142, R298 and Q299 (Gao et al., 2023). In the dimer structure, the N-finger of one monomer is squeezed between Domain II and Domain III of the other monomer. This allows residues S1 and R4 of one M^pro^ monomer to interact with E166 and E290 of the other monomer, respectively, adjusting the orientation of the S1 pocket of the active site for substrate binding (Sacco et al., 2020). R4A and E290A were previously reported to eliminate activity for SARS-CoV M^pro^ (Chou et al., 2004). Similarly, M^pro^ carrying E166A showed markedly lower enzymatic activity (Jochmans et al., 2023).

**Figure 1.**
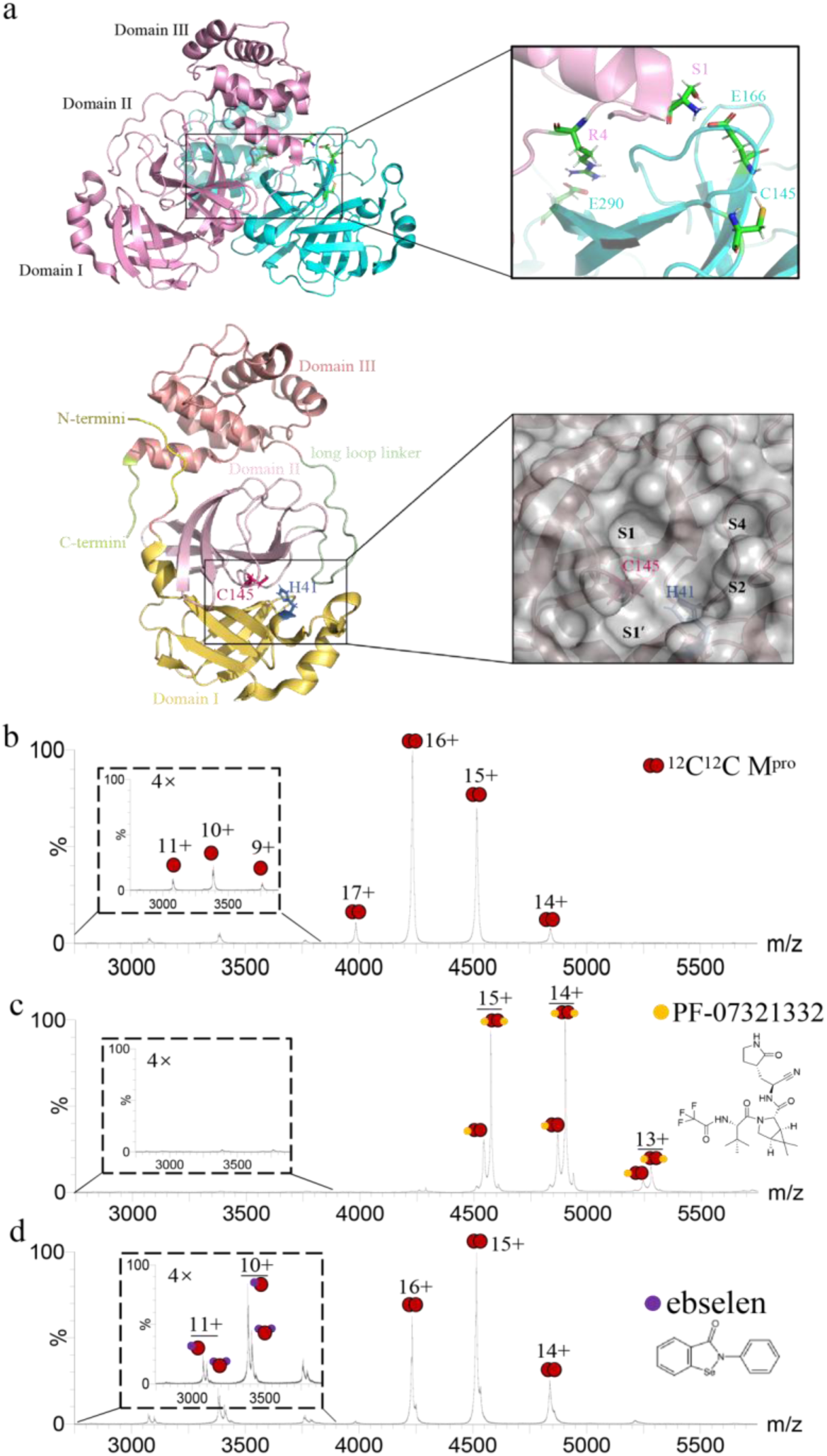
The monomer-dimer equilibrium modulation of M^pro^ induced by inhibitor binding. (a) Crystal structure of SARS-CoV-2 M^pro^ dimer, monomer, and active site pockets of SARS-CoV-2 M^pro^ with subsites and two catalytic dyad residues (H41 and C145) highlighted (PDB entry 7ALI). The upper structure represents the dimer, and the lower structure represents the monomer, respectively. Native mass spectra of (b) 2.5 μM unbound M^pro^, (c) M^pro^-PF-07321332 complex (2.5 μM M^pro^ with 3.75 μM PF-07321332), and (d) M^pro^-ebselen complex (2.5 μM M^pro^ with 7.5 μM ebselen).

Various M^pro^ inhibitors have been developed to target M^pro^, primarily by repurposing clinically approved drugs or adapting inhibitors originally designed against SARS-CoV M^pro^ (Hoffman et al., 2020; Yang and Yang, 2021). Among these, many inhibitors are peptidomimetic compounds that covalently bind to C145 within the active site, such as PF-07321332, PF-00835231, GC376, and boceprevir (Table 1). PF-07321332 (nirmatrelvir) is the first orally administered inhibitor targeting SARS-CoV-2 M^pro^, showing a significantly low EC_50_ (0.075 μM) (Zhao et al., 2022). PF-00835231, initially designed for the M^pro^ of SARS-CoV, also inhibits SARS-CoV-2 with a low EC_50_ of 0.23 μM (Owen et al., 2021). GC376, a known cysteine protease covalent inhibitor targeting multiple viruses, exhibited antiviral activity against SARS-CoV-2 with an EC_50_ of 0.92 μM (Bai et al., 2021). Boceprevir, an approved serine protease inhibitor for treating Hepatitis C virus, demonstrated an EC_50_ of 15.57 μM against SARS-CoV-2 (Fu et al., 2020). Non-peptidomimetic inhibitors targeting the M^pro^ active site have also been identified, e.g., carmofur, ebselen, and MR6-31-2. Carmofur, an antineoplastic agent for colorectal cancer, exhibited an EC_50_ of 24.3 μM against SARS-CoV-2 (Jin et al., 2020b). Ebselen, an organoselenium molecule, forms a seleno-sulfide bond with thiol groups of cysteine on various proteins, leading to anti-inflammatory, antimicrobial, and neuroprotective effects (Azad and Tomar, 2014). Ebselen exhibits low cytotoxicity, and its safety profile has been assessed in multiple human clinical trials (Kil et al., 2017; Lynch and Kil, 2009; Masaki et al., 2016). Ebselen and its derivative MR6-31-2 displayed promising anti-SARS-CoV-2 activity with EC_50_ values of 4.67 μM and 1.78 μM, respectively (Amporndanai et al., 2021). The lower EC_50_ of MR6-31-2 may reflect differences in compound stability, cellular permeability, intracellular exposure, or M^pro^ binding, arisen from its modified chemical structure. However, cell-based EC_50_ values do not directly represent biochemical inhibitory potency against purified M^pro^. While these inhibitors mentioned above all covalently bind to C145, they occupied different subsites (Figure S1) and formed different hydrogen bonds with M^pro^ (Table 1), possibly correlating with their distinct inhibitory effect. Allosteric inhibitors that non-covalently bind outside the active site of SARS-CoV-2 M^pro^ have been characterized via X-ray screening (Günther et al., 2021). One such allosteric site is located in the cavity between the catalytic and dimerization domains, to which AT7519 binds with an EC_50_ of 25.16 μM. Additionally, the C-terminal dimerization domain forms a hydrophobic pocket, which accommodates the aromatic moiety of molecules, exemplified by pelitinib, which exhibits an EC_50_ of 1.25 μM.

**Table 1.**
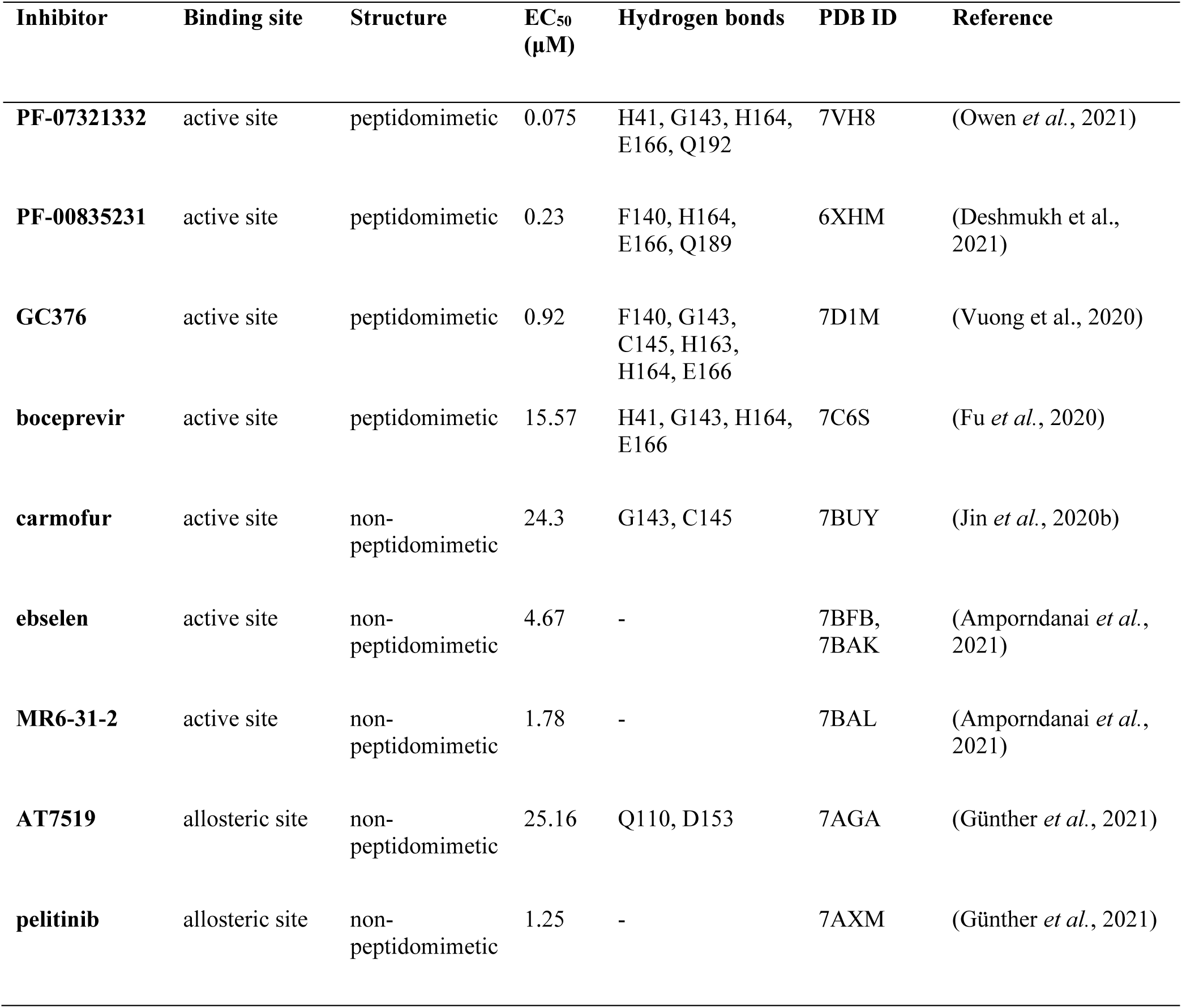
Inhibitors against SARS-CoV-2 M^pro^ and their inhibitory properties.

While these seven covalent-binding inhibitors showed marked differences in inhibitory effects (EC_50_, Table 1), structural alignment of M^pro^ showed that the active site pocket does not undergo significant structural changes upon inhibitor binding (Figure S2). This suggests that the divergent inhibitory effects are not attributable to major structural changes of M^pro^ but likely arise from conformational changes of M^pro^ induced by inhibitor binding. Hydrogen/deuterium exchange mass spectrometry (HDX-MS) is a powerful technique for studying conformational dynamics of proteins and protein-drug complexes in solution by monitoring deuterium exchange of the amide hydrogens of protein backbone (Huang et al., 2019; James et al., 2021; Kish et al., 2023; Narang et al., 2020; Stofella et al., 2024). Typically, amide hydrogens in more exposed and/or flexible regions exchange faster than those at buried and/or rigid regions.

Therefore, alterations in protein structures and dynamics could be reflected by changes in HDX behaviors of amide hydrogens. Our group has previously applied HDX-MS for studying the conformational dynamics of β-lactamases upon binding with inhibitors, revealing promising allosteric sites as targets of drug development (Huang et al., 2020a; 2021; Huang et al., 2025). Current HDX-MS investigations of M^pro^ remain inadequate for a comprehensive elucidation of the inhibitory effects of these inhibitors, as only the effect of PF-07321332 among these nine inhibitors on conformational dynamics of M^pro^ has been presented by HDX-MS (Greasley et al., 2022; Yadav et al., 2022). Furthermore, the dimerization process of M^pro^, which is essential for its enzymatic function, represents a promising yet relatively underexplored target in the field of drug discovery. Intriguingly, the inhibitor PF-07321332, despite targeting the active site, has been reported to promote M^pro^ dimerization by shifting the monomer-dimer equilibrium toward the dimeric state (Higashi-Kuwata et al., 2023). This implies that the alterations in the dimerization process may be another effect to reflect effective inhibitor binding, other than mainly via direct blockade of the active site. To comprehensively advance our understanding of the inhibitory mechanisms, it is essential to investigate the effects of these inhibitors on the dimerization-related monomer-dimer equilibrium of M^pro^ and elucidate the conformational dynamics alterations of M^pro^ induced by inhibitor binding.

In this study, we investigated the effects of three different types of inhibitors on the monomer-dimer equilibrium and conformational dynamics changes of M^pro^ by integrating isotope labeling, native MS, and HDX-MS. We characterized the significantly opposite effects of peptidomimetic inhibitors and ebselen on monomer-dimer equilibrium and conformational dynamics of M^pro^. These findings underscore the importance of the dimer interface regions as allosteric sites, offering valuable clues for the design of inhibitors targeting M^pro^ dimerization.

## Results

### The monomer-dimer equilibrium modulation of M^pro^ induced by inhibitor binding

M^pro^ was characterized by native MS, and two well-resolved charge state distributions were readily identified. The results revealed an equilibrium in solution between monomer (9+ to 11+) and dimer species (14+ to 17+) of M^pro^, with the dimer being the predominant form (Figure 1b), consistent with reported native MS results of M^pro^ (El-Baba et al., 2020). Deconvolution of the mass spectrum yielded a mass of 33,796 Da for the monomeric form (theoretical mass: 33,797 Da), and 67,840 Da for the dimeric form (theoretical mass: 67,594 Da). The 246 Da mass increment relative to the theoretical prediction of the dimer is likely due to the addition of solvent molecules and buffer ions, presumably at the dimer interfaces (Nettleton et al., 1998). Different inhibitors exert diverse effects on M^pro^ monomer-dimer equilibrium (Figures 1c,d and S3). Quantitative analysis of triplicate native MS measurements showed that monomeric M^pro^ accounted for 5.35 ± 0.15% of the total detected M^pro^ population in the absence of inhibitors. PF-07321332 markedly reduced the monomer fraction to 0.48 ± 0.06%, consistent with stabilization of the dimeric state. The ratio of monomers decreased, and that of the dimer increased upon PF-07321332 binding, suggesting the monomer-dimer equilibrium of M^pro^ shifted toward the dimeric form (Figure 1c). This indicates that PF-07321332 not only binds to M^pro^ but also promotes its dimerization. The other three peptidomimetic inhibitors PF-00835231, GC376, and boceprevir also showed similar trends (Figure S3a-c). The corresponding monomer fractions were 0.74 ± 0.18%, 0.43 ± 0.02%, and 0.42 ± 0.02% for PF-00835231-, GC376-, and boceprevir-bound M^pro^, respectively, all substantially lower than that of unbound M^pro^ (5.35 ± 0.15%). Taken together, these results suggest that when peptidomimetic inhibitors bind to the active site C145, they may induce the conformational change of M^pro^. This perturbation may propagate from the active site to the dimer interface, ultimately promoting dimerization. Intriguingly, the allosteric inhibitor AT7519 also shifted the equilibrium toward the dimeric state (Figure S3f), with the monomer fraction significantly decreasing to 2.23 ± 0.21%, compared with 5.35 ± 0.15% for unbound M^pro^. This observation aligns with previously published molecular dynamics (MD) simulation findings that AT7519 enhances both intra- and inter-domain coupling of M^pro^ (Zhang et al., 2024). Notably, the two non-peptidomimetic inhibitors, carmofur and MR6-31-2, did not significantly alter the monomer-dimer equilibrium (Figure S3d,e). The monomer fractions were 5.12 ± 0.89% and 6.46 ± 0.62% in the presence of carmofur and MR6-31-2, respectively, with neither differing significantly from unbound M^pro^ (5.35 ± 0.15%, p > 0.05). Despite also covalently binding to the active site C145, the impacts of carmofur and MR6-31-2 on the equilibrium differed from those of peptidomimetic inhibitors. The hydrogen bond interactions of residues E166 and S1 are crucial for M^pro^ dimerization (Sacco *et al*., 2020), and E166 can also form hydrogen bonds with all peptidomimetic inhibitors but not with non-peptidomimetic inhibitors (Table 1). The residue S1 from one monomer, the peptidomimetic inhibitor, and E166 from the other monomer form tight interactions, revealed by the enhanced hydrogen bond strength of residues S1 and E166 upon peptidomimetic binding compared to unbound M^pro^ (Figure S4). This bolstering of M^pro^ dimerization induced by peptidomimetic inhibitors may reflect the highly effective inhibitory effect of peptidomimetic inhibitors, as they firmly occupy the enzymatic active site pocket and competitively prevent the entry of SARS-CoV-2 immature polyproteins, thereby effectively preventing viral replication. It is worth noting that the promoted dimerization of the protease induced by inhibitor binding has been observed in the interactions between Human immunodeficiency Virus type 1 (HIV-1) protease and its inhibitors. In the presence of HIV-1 protease peptidomimetic inhibitors such as saquinavir and nelfinavir, levels of HIV-1 protease monomers decreased as they formed dimers bound to the inhibitors (Hayashi et al., 2014). This highlights the enzyme-specific nature of inhibition shared by both HIV-1 protease and SARS- CoV-2 M^pro^.

Binding of ebselen (Figure 1d) and pelitinib (Figure S3g) to M^pro^ induced a significant shift toward the monomeric form, with monomer fractions increasing to 19.91 ± 2.18% and 11.86 ± 1.62%, respectively, indicating dissociation of the dimer. Protein charge-state distributions can be influenced by solution-phase conformation, conformational flexibility, solvent properties, and electrospray droplet charging (Susa et al., 2017). Pelitinib shifted the dimer charge-state distribution toward lower charge states, which might be related to the changes of the protein conformation, solvent property and electrospray droplet charging with the addition of pelitinib. Pelitinib was reported to engage in hydrophobic interactions with C-terminal amino acid residues at the dimer interface (Song et al., 2024). This binding may negatively impact the interactions among dimer interface residues and disrupt the dimer interface. The destabilization of M^pro^ was also revealed by a decrease in the melting temperature (T_m_) of M^pro^ upon binding with ebselen (Ma et al., 2020). The fragment JGY and nanobody NB2B4 were reported to bind in the dimer interface, destabilize the M^pro^ dimer, and showed inhibitory activity with an IC_50_ value of 100 μM and 150 nM, respectively (El-Baba *et al*., 2020; Sun et al., 2022). This suggests that impairment of the M^pro^ dimer form may be associated with the inhibitory effect. While ebselen was generally reported to bind to C145 for its inhibitory effect, our native MS results revealed more than one ebselen binding to M^pro^ monomer and an increase in the amount of M^pro^ monomers upon ebselen binding compared to unbound M^pro^ monomers (Figure 1d), implying the ebselen-bound monomers are incapable of reassociating into dimers. We speculate that ebselen binding may induce instability in the M^pro^ dimer interface, thereby disrupting the dimer. Ebselen has not been previously reported to impair M^pro^ dimerization, suggesting a potentially novel inhibitory mechanism of ebselen.

### Subunit exchange dynamics of M^pro^ tuned by inhibitor binding

Subunit exchange is an effective process for analyzing the dissociation and association of multimeric protein complexes (Yang et al., 2013), which can be monitored by native MS that can retain and study intact proteins and protein complexes in their native states (Aquilina et al., 2005; Sobott et al., 2002; Thangaraj et al., 2019; Uetrecht et al., 2010). As the monomer-dimer equilibrium is reflected by the subunit exchange of the M^pro^ dimer, we employed isotope labeling native MS to investigate the effect of inhibitors on subunit exchange dynamics of M^pro^. The strategy for subunit exchange is depicted in Figure 2a. As SARS-CoV-2 M^pro^ (^12^C^12^C) is mixed with ^13^C-labeled M^pro^ (^13^C^13^C), a new heterodimer (^12^C^13^C) occurs due to dimer dissociation and monomer reassociation. These three dimers with different masses can be monitored by native MS (Figures 2b and S5a,b). In contrast to free M^pro^, the heterodimer M^pro^ signal was undetected after PF-07321332 binding over the course of 30 minutes (Figure 2c). These findings imply that the PF-07321332 binding induces a structural change that enhances the affinity of the dimer interface, thereby preventing dissociation. Recent microcalorimetry experiments have provided additional evidence of strong stabilization of M^pro^ by PF-07321332, as evidenced by a 10°C shift in T_m_ and a corresponding increase of 10 kcal/mol in enthalpy (Paciaroni et al., 2023). In addition to PF-07321332, peptidomimetic inhibitors PF-00835231, GC376, and boceprevir also significantly inhibited subunit exchange, demonstrating their strong binding capacity with M^pro^, although boceprevir exhibited a relatively weaker effect (Figure 2d-f). These findings are consistent with our monomer-dimer equilibrium results and suggest that peptidomimetic inhibitors can alter the monomer-dimer equilibrium by preventing dimer dissociation. Intriguingly, ebselen also inhibited subunit exchange of M^pro^ to a certain extent (Figure 2g). This effect may be attributed to the binding of ebselen to the M^pro^ monomer, preventing its association into dimers and thereby reducing the population of dimers. Carmofur and MR6-31-2 did not change the subunit exchange (Figure S6b,c), consistent with our monomer-dimer equilibrium results. For AT7519 and pelitinib binding, no significant effect on subunit exchange was observed (Figure S6d,e), different from the monomer-dimer equilibrium findings, possibly due to the reduced M^pro^-to-inhibitor ratio (from 1:15 to 1:3). These results suggest that AT7519 and pelitinib exhibit weak binding affinity for M^pro^, and a higher inhibitor- to-M^pro^ ratio will be required for subsequent HDX-MS study.

**Figure 2.**
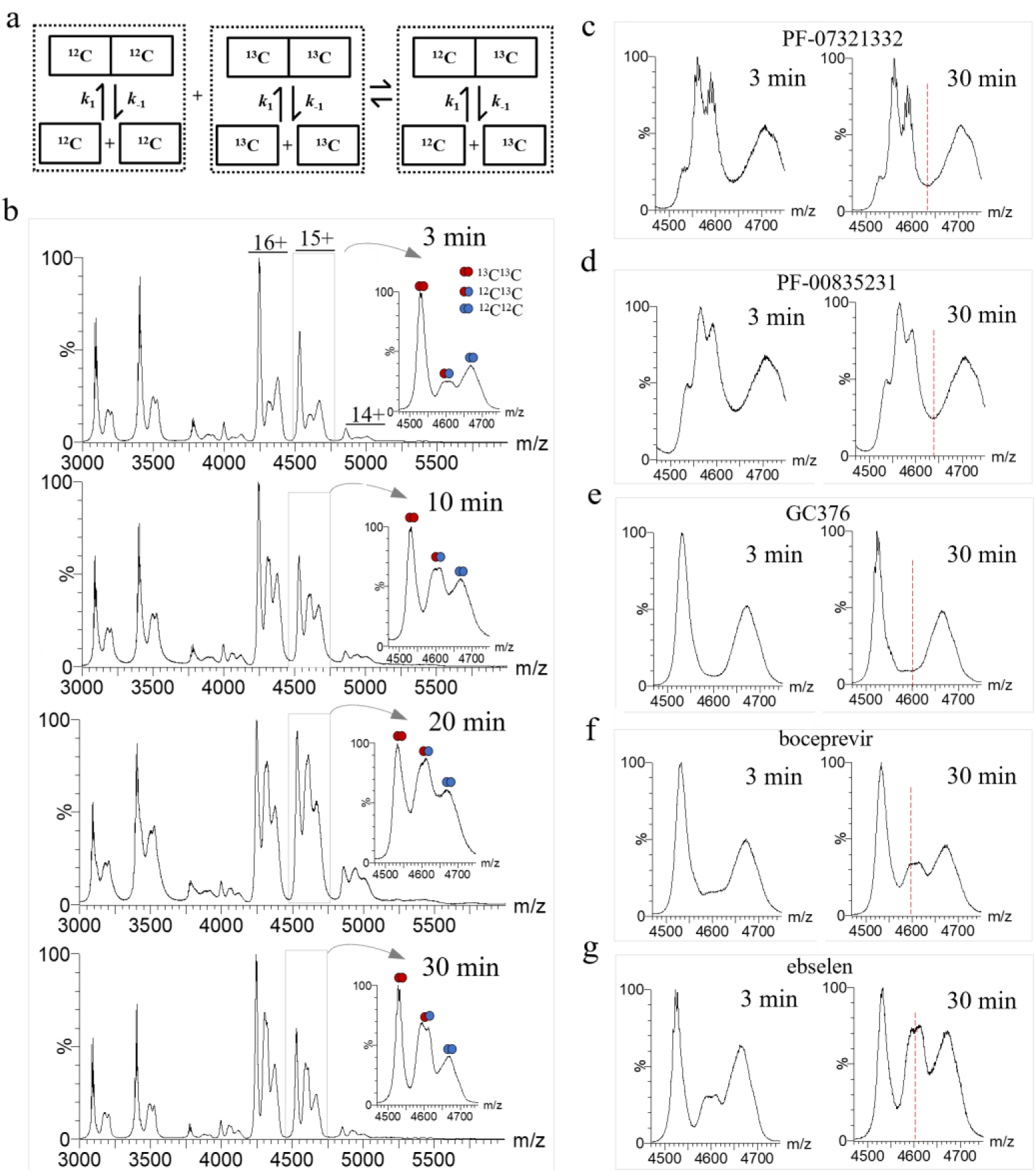
M^pro^ subunit exchange dynamics. (a) The strategy for M^pro^ subunit exchange. (b) Monitoring of unbound M^pro^ subunit exchange by native mass spectrometry. M^pro^ subunit exchange is affected by (c) PF-07321332, (d) PF-00835231, (e) GC376, (f) boceprevir, and (g) ebselen. The red dashed lines indicate the heterodimers.

### Conformational changes of M^pro^ upon inhibitor binding

HDX-MS was employed to provide conformational insights into the binding of inhibitors. By optimizing the injected M^pro^ amount and quenching buffer, pepsin digestion of M^pro^ resulted in 94.4% coverage of the entire sequence (Figure S7). Local HDX-MS of unbound M^pro^ revealed relatively high deuterium uptake in the long loop regions, indicating the relatively high flexibility (Figure 3a). This structural flexibility suggests that the long loop linker may be flexible enough to effectively accommodate different groups of inhibitors. Conversely, the N-terminal residues showed relatively low deuterium uptake, indicating that this region is relatively more rigid (Figure 3a), possibly due to the hydrogen bonds associated with the N-finger.

**Figure 3.**
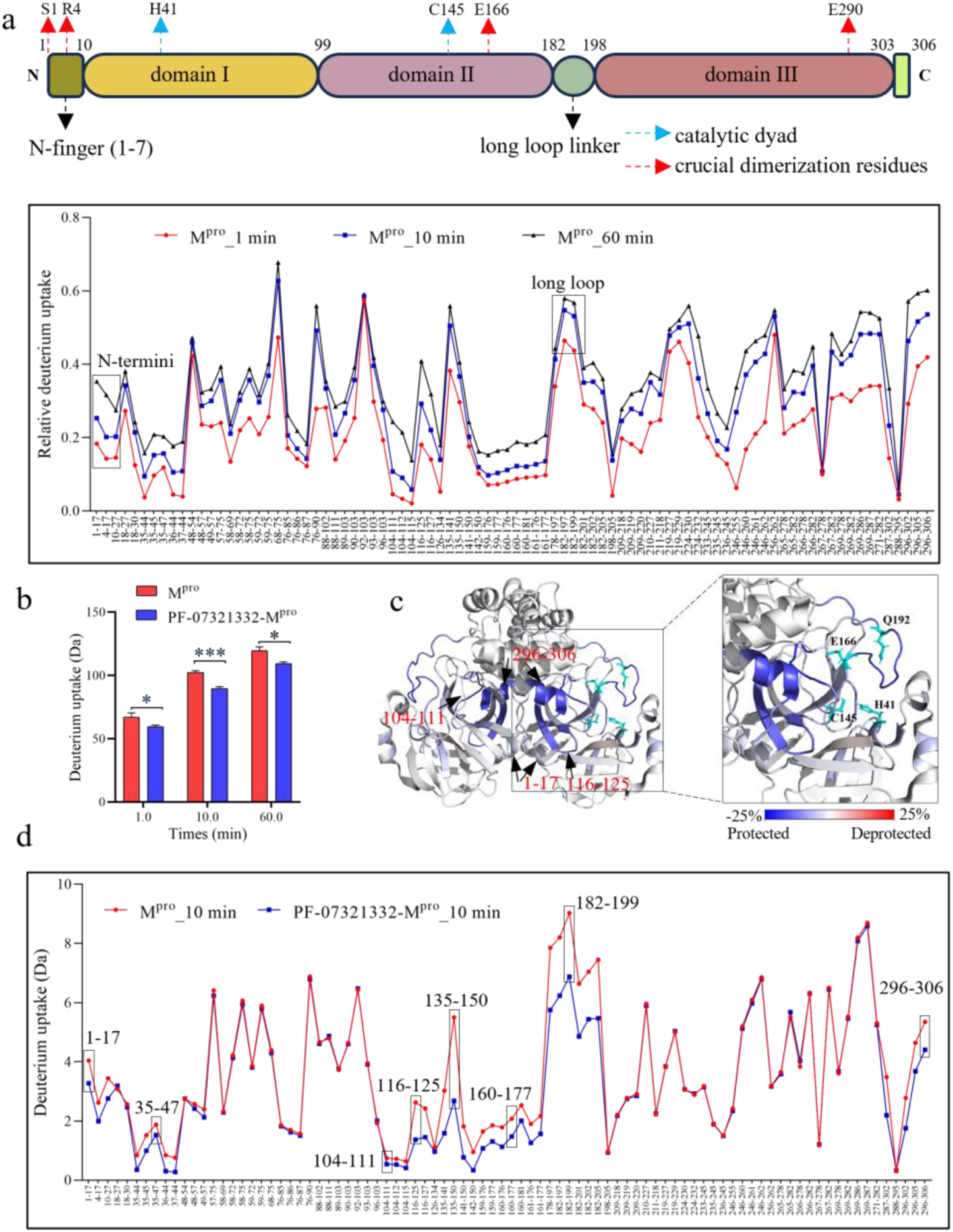
Conformational dynamics analysis of SARS-CoV-2 M^pro^ with PF-07321332. (a) The deuterium uptake plot for all identified peptides from M^pro^ at 1, 10, and 60 min. (b) Deuterium uptake of PF-07321332-bound M^pro^ by global HDX-MS. (c) The heat map of M^pro^ upon binding with PF-07321332 at 10 minutes was overlaid onto the crystal model of M^pro^ (PDB entry 7ALI). (d) The uptake line plots in the PF-07321332-bound M^pro^ at 10-min HDX labelling. Residues 1-17 (***), 35-47 (***), 104-111 (**), 116-125 (***), 135-150 (***), 160-177 (***), 182-199 (***) and 296-306 (***). ** for P < 0.01, and *** for P < 0.001.

The effects of inhibitors on the conformational dynamics of M^pro^ were detected using global and local HDX-MS. The global deuterium uptake of M^pro^ significantly decreased upon binding with PF-07321332, indicating that the overall conformation of M^pro^ became more rigid after the binding (Figure 3b). To obtain more detailed insights into the regional conformational changes of M^pro^, local HDX-MS was performed. Upon binding to PF-07321332, significant changes in HDX were observed in certain active site regions (Figures 3c,d and S9a). Residues 35-47 and 135-150, encompassing the interacting residue H41 and covalent binding site C145, exhibited significant decreases in deuterium uptakes, indicating reduced flexibility, consistent with X-ray crystallography results that H41 and C145 formed strong hydrogen bond and covalent bond interactions with the inhibitor molecule (Zhao *et al*., 2022). Because substitution of the catalytic residue C145, such as C145A, abolishes M^pro^ proteolytic activity, enzymatic assays using this mutant cannot readily distinguish the contribution of C145 to inhibitor binding from its essential catalytic role (Bhandari et al., 2025). Therefore, the effects of PF-07321332 observed here were interpreted together with its established structural and biochemical binding mechanism. Additionally, residues 160-177 in Domain II exhibited reduced deuterium uptake upon PF-07321332 binding, which may be attributed to the hydrogen bond formation between residue E166 and the inhibitor (Table 1). E166 is crucial for the correct orientation of the active site pocket. Therefore, the reduced flexibility of this region could potentially influence the formation of this subsite and consequently suppress the protein’s activity. Interestingly, we found that residues 104-111 and 116-125, which do not directly participate in the inhibitor binding, also showed decreased deuterium uptake and thus reduced flexibility after the binding, suggesting that these regions might modulate the binding of the inhibitor to the active site by a short-range communication, enabling proper conformation. Furthermore, residues 182-199, corresponding to the long loop linker connecting Domain II and Domain III, demonstrated a highly significant decrease in deuterium uptake, implying that the flexibility of this loop was reduced to a large extent upon binding to the inhibitor (Figure 3d). Q192 in this loop forms a hydrogen bond with the inhibitor molecule, which may be the primary interaction responsible for the loop’s lower flexibility upon PF-07321332 binding. The residue D187 in this loop plays a crucial role in maintaining the charge of the catalytic H41 (Kneller et al., 2020). Thus, disturbing the plasticity of this loop upon inhibitor binding could impair the capacity of M^pro^ to accommodate various ligands and hydrolyze substrates, ultimately decreasing its activity. This observation aligns with previously reported M^pro^-inhibitor complex simulations, which suggest that this loop exhibits reduced mobility upon binding to the inhibitor (Suárez and Díaz,

2020). Designing molecules that can interact with this large loop could be an attractive strategy for inhibition. In addition to the findings mentioned above, we newly discovered that the conformational dynamics of the N- and C-termini, which were not directly involved in inhibitor interactions, were significantly affected upon PF-07321332 binding. Residues 1-17 and 296- 306 exhibited notable decreases in deuterium uptake upon the inhibitor binding, indicating that these regions became more rigid. The N- and C-termini are known to be dimer interface residues and crucial for the formation of the active dimeric form (Pekel et al., 2022; Zhao *et al*., 2022). Thus, changes in their conformational dynamics could potentially impact the dimerization process. The conformational rigidity of the termini directly inhibits dimer dissociation, thereby accounting for the shift of the monomer-dimer equilibrium toward the dimeric state.

Comparative HDX-MS analysis revealed distinct conformational dynamics of M^pro^ induced by inhibitor binding. While GC376 and PF-00835231 induced similar HDX changes as those observed with PF-07321332 (Figures S8a,b, S9b,c and S11a,b), the magnitude of deuterium uptake alterations diverged significantly for boceprevir and carmofur compared to PF-07321332 (Figures S8c,d, S9d, S10a and S11c,d). Notably, boceprevir binding induced no significant HDX changes in the N-terminal region, while carmofur binding left both the N- and C-termini uptakes unchanged. These differential HDX profiles correspond with our findings on subunit exchange: boceprevir obviously but incompletely disrupted subunit exchange, whereas carmofur showed no significant effect. Additionally, the active site region showed lower HDX reduction for boceprevir and carmofur relative to PF-07321332, with similar trends observed in residues 104-111, 116-125, and the long loop linker, possibly due to the different binding affinities of the inhibitors for M^pro^. Ligands with higher affinity are generally with slower off-rates, thereby maintaining their interactions with M^pro^ for longer periods and inducing greater protection. Thus, the variations in deuterium uptake likely reflect the differing abilities of inhibitors to stabilize M^pro^ dimerization state, which are influenced by their binding strengths and relative on- and off-rates.

AT7519 and pelitinib are allosteric inhibitors that non-covalently bind outside of the active site of M^pro^ (Günther *et al*., 2021), indicating that M^pro^ may exhibit different conformational dynamics when binding these inhibitors compared to those targeting the active site. Upon binding with AT7519 or pelitinib, no significant differences in deuterium uptake of M^pro^ were observed, even at a 50× molar excess of inhibitor relative to M^pro^ (Figure S10d,e). A similar result was reported, showing that M^pro^ exhibited no significant deuterium uptake difference when binding with pelitinib at a 50:1 molar ratio (pelitinib: M^pro^) (Yadav *et al*., 2022). The lack of observed HDX might stem from experimental limitations in studying the weak interactions of binders using HDX-MS. The non-covalent interactions of AT7519 and pelitinib with M^pro^ might not induce significant perturbations in backbone dynamics that are detectable at the peptide level by HDX-MS.

In contrast, covalent inhibitor ebselen induced distinct conformational dynamics in M^pro^, as global HDX-MS analysis revealed higher structural flexibility upon binding of one and two ebselen molecules, respectively (Figure 4a). This contrasts with the typical covalent inhibitors of M^pro^, suggesting that ebselen may employ an entirely different inhibitory mechanism from others. Ebselen binding resulted in an increased deuterium uptake across most regions of the M^pro^, especially the N- and C-termini (Figures 4b,c and S10b). Our MD simulation also supports this finding (Figure S13b-d), with an increased RMSF after binding. The increased flexibility of these two terminal regions, combined with the native MS findings, indicates that the conformational change of the N- and C-termini may relate to the monomer-dimer equilibrium shift. The deprotection in the N-terminal region was also observed in MR6-31-2 binding, implying a similar inhibitory mechanism to ebselen (Figures S8e, S10c and S11e).

**Figure 4.**
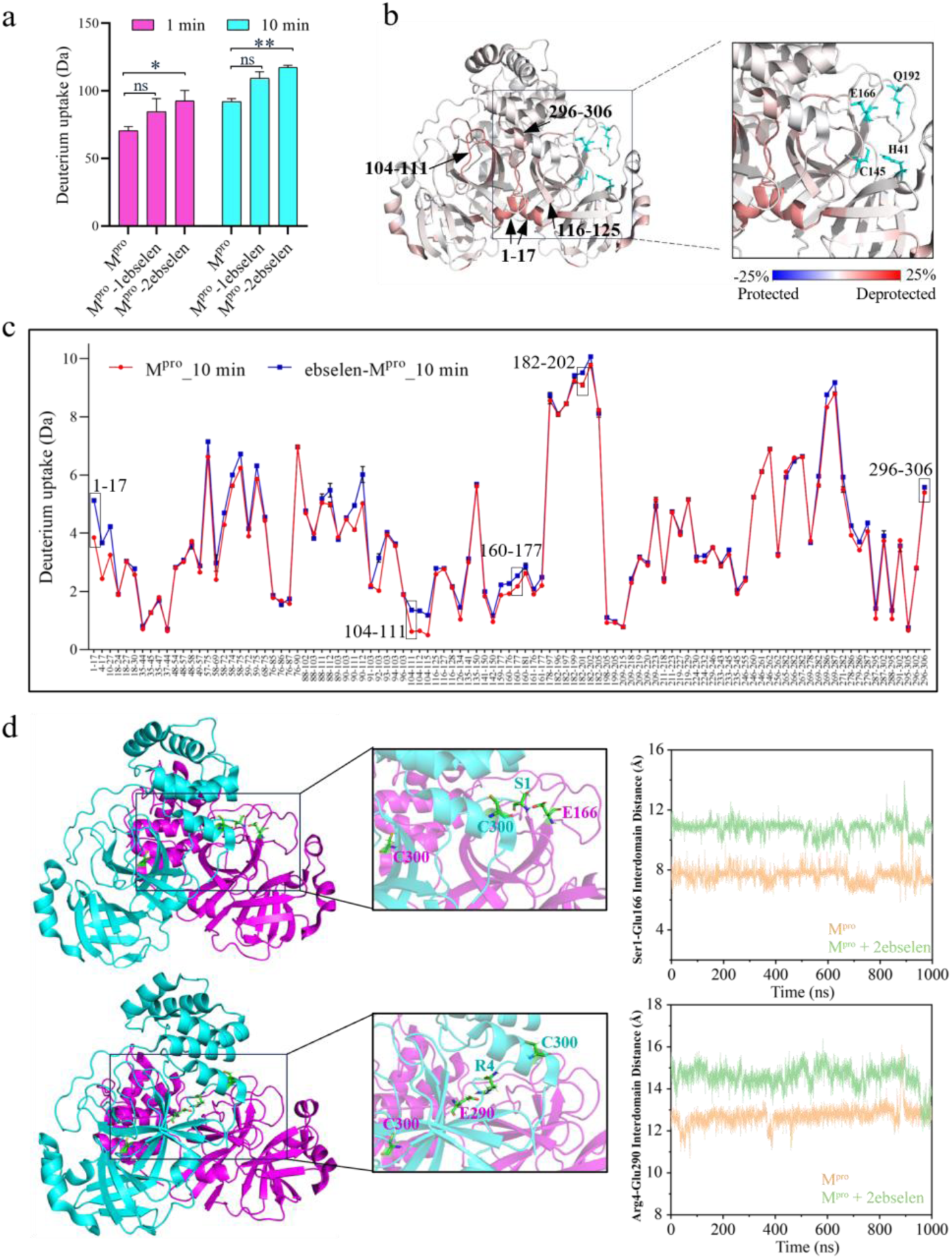
A new allosteric binding mode of ebselen on M^pro^. (a) Deuterium uptake plot of ebselen-bound M^pro^ by global HDX-MS. (b) The heat map of M^pro^ upon binding with ebselen at 10 minutes was overlaid onto the crystal model of M^pro^ (PDB entry 7ALI). (c) The uptake line plots in the ebselen-bound M^pro^ at 10 minutes HDX labelling. Residues 1-17 (***), 104-111 (**), 160-177 (**), 182-202 (**) and 296-306 (*). * for P < 0.05, ** for P < 0.01, and *** for P < 0.001. (d) MD simulation of SARS-CoV-2 M^pro^ in the presence of ebselen on C300. Shifts in key intra-dimer interactions before and after covalent binding of ebselen at C300 on M^pro^. Left panel: key amino acids involved in intra-dimer interactions are depicted on the crystal model of M^pro^ (PDB entry 7ALI). Right panel: the distances between R4 of one monomer and E290 of the other monomer Cα atoms, and residues S1 of one monomer and the other monomer E166 Cα atoms were measured, respectively.

### A new binding site of ebselen on M^pro^

Ebselen revealed multiple M^pro^ binding sites and exhibited distinct conformational dynamics compared to other inhibitors, suggesting it may alter conformational dynamics of M^pro^ by binding to cysteines outside the active site C145. To identify the binding sites of ebselen on M^pro^, liquid chromatography-tandem mass spectrometry (LC-MS/MS) was employed. Incubation of M^pro^ with ebselen revealed ebselen-mediated cysteine modifications at C44 and C300 (Figure S12). However, no adduct was observed on C145, aligning with observations reported in previous structural analysis that a selenium-containing adduct on C145 was identified arising from the hydrolysis reaction of ebselen (Amporndanai *et al*., 2021). However, the selenium-containing peptide with a single selenium atom bound to Cys145 was also not identified in the software-based analysis. This selenium-modified peptide could potentially cross-link with another selenium-containing peptide, which is typically challenging to detect using search software. The crystal structure of the ebselen-M^pro^ complex demonstrates that ebselen forms a covalent bond with C44 (PDB: 7BFB), aligning with our MS findings. The C44A mutation has been shown to inhibit the activity of M^pro^, highlighting a significant role for C44 in M^pro^ activity (Iacobucci et al., 2025). These findings indicate that ebselen may inactivate SARS-CoV-2 M^pro^ by covalently modifying C44, thereby impairing the function of this essential viral protease. Additionally, reported MD simulations results suggested that ebselen can bind at two probable sites: One at C145 within the catalytic cavity via a Se-S bond, and the other at the dimerization region between Domain II and Domain III (Menéndez et al., 2020). M^pro^ could be reversibly inhibited through the oxidation of C300 with glutathione, causing M^pro^ to form an inactive monomer (Davis et al., 2021). Tixocortol has also been identified as an M^pro^ inhibitor that primarily targets C300, impairing M^pro^ dimerization (Davis et al., 2024). Bismuth drug colloidal bismuth subcitrate (CBS) was reported to bind to C300 of M^pro^ and result in dissociation of M^pro^ dimer and proteolytic dysfunction (Tao et al., 2021). These results highlight an unexpected reactivity of additional cysteine residues beyond the catalytic site C145, suggesting that a combination of inhibitory mechanisms of inhibitors, rather than solely blocking the active site, is involved. Our HDX results showed that residues 296-306 in the C-terminal region of M^pro^ were more flexible upon ebselen binding (Figure 4c).

To gain deeper insight into the molecular mechanism of action of ebselen on M^pro^, we conducted an all-atom MD simulation to compare the structures and flexibility of M^pro^ before and after ebselen binding at C300. We performed 1000 ns of MD simulations in triplicate (Figure S13) to investigate the interactions of key amino acids at the dimeric interface (residues S1 and E166, and R4 and E290), which are essential for maintaining the dimeric form of M^pro^. For the ebselen-modified M^pro^, the Cα distance between R4 of the N-finger of one monomer and the E290 of Domain III of the other monomer changed from 7.5 Å to 10.5 Å after ebselen binding. Additionally, the Cα atoms distance between S1 on the N-finger of each monomer and E166 on Domain II of the other monomer increased from 13.0 Å to 15.5 Å (Figure 4d). These observations suggested that covalent modification of ebselen at C300 may reduce the intensity of M^pro^ dimer hydrogen bond, thereby inducing a steric blocking of dimer formation. Given its unique positioning at the dimeric surface, mutation or modification of the conserved C300 may impair M^pro^ dimerization and further cause the active enzyme to form an inactive monomer, offering new avenues for designing novel agents to combat SARS-CoV-2.

Although tandem MS and MD simulations suggested that C300 modification contributed to ebselen-mediated destabilization of the M^pro^ dimer, the functional significance of C300 remained to be experimentally validated. To address this question, C300S and C300F mutants were generated to determine whether elimination of the C300 thiol group influences the intrinsic monomer-dimer equilibrium of M^pro^ and attenuates ebselen-induced conformational changes and functional inhibition. C300S was designed to remove the reactive thiol group while minimally affecting M^pro^ structure and dimerization. C300F was introduced to mimic the steric perturbation caused by ebselen modification and evaluate its impact on M^pro^ dimerization. To evaluate whether C300 mutations affect M^pro^ folding and dimerization, WT M^pro^ and the C300S and C300F variants were analyzed by SDS-PAGE and native PAGE (Figure S14). Under denaturing SDS-PAGE conditions, all three proteins showed major bands at similar apparent masses corresponding to monomeric M^pro^, indicating successful purification without obvious degradation. Native PAGE revealed that WT and C300S M^pro^ predominantly existed as dimeric species, whereas C300F migrated mainly as a monomeric species. These results indicate that the C300S substitution preserves the dimeric architecture of M^pro^, whereas C300F markedly impairs dimer formation. The differential effects are likely due to the relatively conservative nature of the C300S substitution, as serine is small and polar and can partially preserve local packing, whereas the bulky aromatic phenylalanine side chain may introduce steric hindrance within the C-terminal dimer-interface region. Thus, C300S provides a suitable mutant for further evaluating the contribution of C300 thiol modification to ebselen-mediated perturbations, as it preserves the overall dimeric architecture of M^pro^ while eliminating the reactive thiol group.

To determine whether C300 contributes to ebselen-induced perturbation of M^pro^ dimerization, we analyzed C300S M^pro^ in the absence and presence of ebselen by native MS (Figure S15). C300S M^pro^ was incubated with ebselen at a molar ratio of 1:3. Ebselen-bound species were detected for both monomeric and dimeric C300S M^pro^, indicating that ebselen can still bind to the C300S mutant through C300-independent sites. However, unlike WT M^pro^, ebselen did not markedly shift the monomer–dimer equilibrium of C300S toward the monomeric state. These results suggested that C300 is a key residue mediating the ebselen-induced disruption of M^pro^ dimerization. Nevertheless, the persistence of ebselen binding to C300S indicated that ebselen might also interact with additional cysteine residues, such as C145, which could contribute to direct enzymatic inhibition. Therefore, enzymatic activity assays were performed to evaluate the functional consequences of ebselen binding to WT and C300S M^pro^.

We first determined whether the C300 substitutions themselves affected proteolytic function. Relative to WT M^pro^, C300S retained approximately 70% activity, whereas C300F retained only approximately 10% (Figure S16a). The severe activity loss of C300F is consistent with its predominantly monomeric behavior in native PAGE and supports the requirement for an intact C-terminal dimer interface in formation of the catalytically competent enzyme. By contrast, the more conservative C300S substitution preserved both substantial activity and dimer formation. Although its basal activity was moderately reduced, C300S remained sufficiently functional to serve as an informative control for evaluating the specific contribution of the C300 thiol to ebselen inhibition. We therefore compared ebselen inhibition of WT and C300S M^pro^ after normalizing each treated sample to its corresponding untreated control. Ebselen reduced WT activity to approximately 53%, corresponding to approximately 47% inhibition. Under the same conditions, C300S retained approximately 78% activity and showed only approximately 22% inhibition (Figure S16b). The C300S substitution therefore significantly attenuated the functional effect of ebselen. Together with the native MS results, these data provide complementary evidence that C300 modification contributes to ebselen-induced destabilization of the active M^pro^ dimer and to the associated loss of proteolytic activity. The residual inhibition of C300S further indicated that C300 is not the sole functional target of ebselen and is consistent with a multi-site mechanism involving additional cysteine residues, including residues within or near the catalytic region. Overall, the mutagenesis results experimentally supported the model derived from MS/MS, HDX-MS, and MD simulations: ebselen combines C300-dependent allosteric disruption of dimerization with C300-independent inhibition at other reactive sites.

## Discussion

This study investigates SARS-CoV-2 M^pro^ as a critical antiviral drug target and reveals the distinct effects of peptidomimetic inhibitors and ebselen on SARS-CoV-2 M^pro^ dimerization and conformational change. Peptidomimetic inhibitors stabilize the M^pro^ dimer form by rigidifying the N- and C-termini, thereby suppressing subunit exchange of M^pro^ dimer and shifting the monomer-dimer equilibrium of M^pro^ toward the dimeric state. The interaction of M^pro^ and peptidomimetic may firmly lock dimeric M^pro^ in a compact M^pro^-inhibitor complex state, which is inactive for M^pro^ proteolytic activity by effectively competing with substrates for binding and inhibiting substrate cleavage. In contrast, ebselen covalently binds to C300, a critical residue at the dimer interface, thereby inducing conformational flexibility in the C-terminal region and shifting the monomer-dimer equilibrium of M^pro^ toward the inactive monomeric state, representing a novel allosteric inhibition mechanism of ebselen. The conserved nature of the M^pro^ dimer interface across coronavirus species suggests that targeting this region could lead to the development of broad-spectrum antivirals. Especially, the identification of C300 as a druggable allosteric hotspot offers an alternative strategy to combat drug resistance arising from active site mutations.

More generally, the therapeutic potential of inhibitors can be readily assessed by the relatively straightforward dual MS approaches, which involve probing the monomer-dimer equilibrium and the conformational dynamics of M^pro^ upon inhibitor binding. These measurements elucidate the mechanism of action of these inhibitors, either through binding to the active site or by influencing the dimer interface, thereby disrupting or favoring dimerization. This work establishes a paradigm for exploring allosteric modulation in SARS-CoV-2 M^pro^ and provides new ways for assessing the efficacy of novel antiviral drug candidates against M^pro^ in SARS-CoV-2 and emerging coronaviruses.

## Methods

### Materials

PF-07321332, PF-00835231, boceprevir, carmofur, ebselen, AT7519, and pelitinib were purchased from MedChemExpress (Princeton, NJ, USA). GC376 was purchased from Selleckchem (Houston, USA). MR6-31-2 was purchased from Sigma-Aldrich (St. Louis, USA). All inhibitors were dissolved in 100% dimethyl sulfoxide (DMSO) to achieve the desired concentrations, while ensuring that the DMSO concentration remained below 5% in all inhibitor-protein mixtures. Tag-free authentic SARS-CoV-2 M^pro^ (Cat# SAE0172) was from Sigma-Aldrich (St. Louis, USA). Tag-free SARS-CoV-2 M^pro^, C300S M^pro^, C300F M^pro^ and ^13^C-labeled SARS-CoV-2 M^pro^ were produced from Beijing Anbiqi Biotechnology Co., Ltd (Beijing, China). Deuterium oxide was purchased from Cambridge Isotope Laboratories (Tewksbury, USA). Formic acid (MS grade) was purchased from Sigma-Aldrich (St. Louis, USA), and acetonitrile (LC-MS grade) was acquired from Anaqua (DE, USA). Ultra-centrifugal filters with 10 kDa molecular weight cut-off were purchased from Millipore (Burlington, USA).

### Native mass spectrometry

Native MS analysis was performed using nano-electrospray ionization (nano-ESI) on a Waters Synapt G2-Si. The protein samples were buffer-exchanged into 200 mM ammonium acetate utilizing 10 kDa cut-off ultra-centrifugal filters. M^pro^ was incubated with inhibitors for 30 minutes at room temperature before ESI. Typically, 3 μL sample was loaded into homemade gold-coated glass nano-ESI emitters for analysis. Nano-ESI capillaries were fabricated in-house from borosilicate glass tubes with outer and inner diameters of 1 mm and 0.75 mm, respectively (Sutter Instruments, Hercules, CA). Preparation involved using a P-2000 laser-based micropipette puller (Sutter Instruments, Hercules, CA), followed by gold coating with a 150R S Plus sputter coater (Electron Microscopy Sciences, PA, USA). Typical settings in positive ion mode included: capillary voltage at 1.5 kV, sampling cone at 30 V, source offset at 30 V, source temperatures at 30℃, gas flows cone gas at 30 L/h, trap collision energy (CE) at 50 V, transfer CE at 50 V, and gas control trap at 6 mL/min. All spectra were internally calibrated using a cesium iodide solution, with spectral acquisition over a range of 1000-7000 *m/z*. Data analysis was performed using MassLynx v4.2 software, with no background subtraction and minimal smoothing applied.

### Subunit exchange

Label-free SARS-CoV-2 M^pro^ was mixed with ^13^C-labeled SARS-CoV-2 M^pro^ for varying time intervals, and the extent of subunit exchange was monitored using native MS. To investigate the effect of inhibitors on the M^pro^ subunit exchange, M^pro^ was incubated with inhibitors at a ratio of 1:3 (5 μM:15 μM) for 30 minutes at room temperature before mixing with an equal volume of 5 μM ^13^C-labeled SARS-CoV-2 M^pro^. The extent of the subunit exchange was monitored at 3 min and 30 min, respectively.

### Global and local Hydrogen/deuterium exchange mass spectrometry (HDX-MS)

HDX-MS experiments were performed on a Waters Synapt G2-Si, following protocols similar to those described in our previous studies (Huang et al., 2020b). M^pro^ was incubated with covalent inhibitors at a ratio of 1:3 or allosteric inhibitors at a ratio of 1:50 for 30 minutes at room temperature before HDX experiments. 10 pmol of samples (for global HDX-MS) or 50 pmol of samples (for local HDX-MS) were labeled with 9× volume deuterated PBS buffer at various time points. Each sample was then quenched by diluting 1:1 into a quenching buffer (3.2% formic acid in ddH_2_O). For global HDX-MS analysis, the quenched mixtures were loaded onto a Waters ACQUITY UPLC Protein BEH C4 VanGuard Pre-column (300 Å, 1.7 µm, 2.1 mm × 50 mm) and washed for 3 minutes. Subsequently, samples were separated on a Waters ACQUITY UPLC Protein BEH C4 Column (300 Å, 1.7 µm, 2.1 mm × 100 mm), using a 10-minute linear gradient from 20% to 80% solvent B (0.1% FA, 100% acetonitrile). For local HDX-MS analysis, the quenched mixture underwent online pepsin digestion at 20°C using a Water BEH Enzymate Pepsin Column (300 Å, 5 µm, 2.1 mm × 30 mm). The resulting peptides were trapped on a Waters ACQUITY UPLC BEH C18 VanGuard Pre-column (130 Å, 1.7 µm, 2.1 mm × 5 mm) and desalted for 3 minutes. Subsequently, the peptides were separated on a Waters ACQUITY UPLC BEH C18 Column (130 Å, 1.7 µm, 1 mm × 100 mm), using a 12-minute linear gradient from 5% to 95% solvent B. For peptide identification, non-deuterated samples were analyzed using the same LC method. The reference mass of each peptide was generated using the same HDX-MS procedure, in which all D_2_O was replaced with ddH_2_O. A clean blank was injected between each analytical run to eliminate any carryover.

### HDX-MS data analysis

Data for each time point were collected in triplicate. The results are presented as mean values ± standard deviation (SD), where error bars indicate the standard deviation for each time point. For global HDX-MS data, the acquired spectra were deconvolved using MassLynx 4.2 to determine the deuterated protein mass at various time points. Deuterium uptake was calculated by subtracting the mass of the non-deuterated protein from that of the deuterated protein using Microsoft Excel. Pairwise comparisons of deuterium uptake between different states were statistically analyzed using a two-tailed Student’s *t*-test. Significance levels are denoted based on the *t*-test results as follows: *ns* for *P* > 0.05 (not significant), * for *P* < 0.05, ** for *P* < 0.01, and *** for *P* < 0.001. For local HDX-MS data, the MS^E^ spectra were processed with ProteinLynx Global Server 3.0.2 (PLGS, Waters, UK) for peptide identification. The resulting peptides were further filtered using DynamX 3.0 (Waters, UK) with the following parameters: a minimum intensity of 5000, a minimum product ions per amino acid of 0.15, a maximum ppm mass error on the precursor ion of 10, and a minimum score of 6.5. HDX measurements were performed in triplicate, and the relative deuterium uptake for peptides was based on the average of these replicates. Sequence coverage, redundancy, number of generated peptides, repeatability, and significant difference in HDX were calculated automatically using Deuteros software (Lau et al., 2021). A significance threshold was established to assess differences in HDX between states compared in each dataset, corresponding to a 99% confidence interval (CI) calculated from triplicate measures for each comparative HDX dataset. A hybrid significance test with a 99.0% confidence interval was employed for all datasets presented in Woods plots using Deuteros 2.0. The line plots and bar plots were generated using GraphPad Prism 8. For visualization, relative fractional deuterium uptake and relative fractional deuterium uptake differences were mapped onto the M^pro^ crystal structure (PDB entry 7ALI) using PyMOL 3.1. To facilitate access to the HDX data obtained in this study, the HDX Summary Table is included in the Supporting Information, as recommended by the community (Masson et al., 2019).

### Identification of binding sites of ebselen on M^pro^

M^pro^ was incubated with ebselen at a molar ratio of 2:1 and 1:3 at room temperature for 30 minutes, respectively. The mixture was then denatured and digested with trypsin. The resulting peptides were desalted and dried. Samples were analyzed on a Bruker timsTOF Pro 2 Mass Spectrometer (Bruker, Bremen, Germany) coupled with a Thermo Fisher Scientific UltiMate 3000 RSLCnano (Waltham, USA). Dried peptide samples were dissolved in 0.1% formic acid in water and loaded onto an Acclaim™ PepMap™ 100 C18 trap column (5 mm × 1 mm; particle size, 5 μm; pore size, 100 Å; Thermo Fisher Scientific, Waltham, USA). Peptides were then separated on an Aurora™ ULTIMATE C18 analytical column (25 cm × 75 μm; particle size, 1.7 μm; pore size, 120 Å; IonOpticks, Fitzroy, Australia) using a gradient of 5–35% mobile phase B (acetonitrile and 0.1% formic acid) at a flow rate of 300 nL/min over 45 minutes. MS and MS/MS spectra were acquired with a mass scan range of 100-1700 *m/z* and an ion mobility scan range of 0.6-1.6 V•s/cm^2^. The dual TIMS setup enabled operation at a 100% duty cycle, with ramp and accumulation times set at 100 ms. MS/MS spectra were acquired using DDA-PASEF (Data-dependent acquisition-Parallel Accumulation-Serial Fragmentation) mode. Database searches were performed using PEAKS Studio 12.0, employing a database downloaded from the NCBI Reference Sequence (YP_009725301.1) for SARS-CoV-2 M^pro^. The protein identification search was conducted with the following parameters: variable modifications included methionine oxidation (+15.995 Da), acetylation (+42.011 Da) at the N-terminus, and cysteine ebselen modification (+275 Da); trypsin digestion was allowed with up to two missed cleavages. Other parameters were set to the software defaults.

### Molecular dynamics (MD) simulation

The initial conformation of the SARS-CoV-2 M^pro^ model was obtained from the PDB database (PDB ID: 7ALI). We also constructed a model of ebselen-bound M^pro^ (to C300, both chains). We referred to another crystal structure (PDB ID: 7BFB). In 7BFB, an ebselen ligand was found near C300 of chain B (not in chain A), but most of the ligand is not defined (has no electron density) except for the selenium. The geometry of the ebselen-modified C300 was poor (S-Se bond too long). Therefore, we built the ebselen-modified M^pro^ model such that it is covalently bound to a suitable rotamer of the C300 residue of both protomers, while avoiding steric clash with nearby protein atoms. The covalent bond between ebselen and C300 is patched with PyMOL and optimized with Sculpting. MD simulations were set up using the Amber 22 software package with GPU-accelerated PMEMD (Particle Mesh Ewald Molecular Dynamics) (Case et al., 2023). The ff14SB force field was applied to generate parameters for the protein (Maier et al., 2015). Because the ligands contain selenium atoms, which are not included in the General Amber Force Field (GAFF), the restrained electrostatic potential (RESP) protocol with HF/6-31G* was employed to calculate the atomic charges for the ebselen. Force field parameters for ebselen and modified cysteine were determined using the Antechamber module of Amber 22. Na(I) counter ions were included to neutralize the charges. The protein structure was solvated in a TIP3P periodic box with a minimum distance of 10 Å from the box edge. The first minimization involves 6000 steepest descent steps with a constant volume periodic boundary, and a second minimization (6000 steps) with no constraints of conjugate gradient energy minimization. Next, the temperature gradually increased from 0 to 298 K over 20 ps using a Langevin thermostat. Density equilibration and production runs were carried out using a constant pressure ensemble (NPT). The SHAKE was applied to constrain the bond length of hydrogen atoms at equilibrium and production stages. Long-range electrostatic interactions were treated using the Particle Mesh Ewald (PME) method, with a nonbonded interaction cutoff of 8 Å. For both the SARS-CoV-2 M^pro^ and SARS-CoV-2 M^pro^-ebselen systems, three independent trajectories were performed with random seeds. Simulations were performed at the same temperature with HDX-MS (298 K) for 1000 ns, with a step size of 200 ps, resulting in a total of 5000 snapshots. Root mean square deviation (RMSD), root mean square fluctuation (RMSF), and distances between specified atoms in each MD snapshot were calculated using the CPPTRAJ module in AMBER 22.

### SDS-PAGE and native PAGE

Protein purity, apparent mass, and oligomeric state were assessed by SDS-PAGE and native PAGE. For SDS-PAGE, 3 μg of wild-type (WT), C300S, or C300F SARS-CoV-2 M^pro^ was mixed with 5× SDS loading buffer, heated at 95 °C for 10 min, and separated on a 10% polyacrylamide gel. For native PAGE, SDS and reducing agents were omitted from the sample buffer, running buffer, and gel to preserve non-covalent oligomeric assemblies. Samples were mixed with 5× non-denaturing loading buffer without heating and separated on a 10% native polyacrylamide gel (Beyotime, Shanghai, China). Gels were stained with Coomassie Brilliant Blue R-250 and destained with ddH_2_O.

### M^pro^ enzymatic activity and inhibition assay

The proteolytic activities of WT, C300S, and C300F M^pro^ were determined using the fluorogenic substrate MCA-AVLQSGFR-Lys(Dnp)-Lys-NH2 (Beyotime, Shanghai, China). Fluorescence was recorded on a Spark multimode microplate reader (Tecan, Männedorf, Switzerland) at excitation and emission wavelengths of 325 and 393 nm, respectively. Basal activity was measured immediately after mixing 0.1 μM M^pro^ with 10 μM substrate. For inhibition assays, M^pro^ was pre-incubated with ebselen for 20 min at room temperature before substrate addition; the final reaction contained 0.1 μM M^pro^, 0.3 μM ebselen, and 10 μM substrate. Reactions were monitored in black 96-well plates at 37 °C for 30 min. Initial rates were obtained by linear regression of the initial linear region of each progress curve. Activities were normalized either to untreated WT M^pro^ or to the corresponding untreated enzyme, as indicated. All measurements were performed in triplicate.

## Supporting information

Source Data of the HDX-MS Studies

## Acknowledgments

We thank Dr. Yu Wai Chen, Dr. Tsz Fung Wong, Ms. Jiayu Lin and other members of the Yao group for their assistances and helpful discussion. We thank Dr. Sirius Pui-Kam Tse, Dr. Pui-Kin So, and Dr. Chi-Hang Chow for their assistance with the project. We thank Dr. Yangyang Sun (Shenzhen Institutes of Advanced Technology, Chinese Academy of Sciences) for the technical support. This work was supported by the National Key Research and Development Program of China (Grant No. 2024YFF0725800), and Hong Kong Research Grants Council (Grant Nos. 15304022, 15308923, R5013-19, C5026-24G, C5031-14E, C4014-23G, and CRS_CUHK405/23). We thank the University Research Facility in Life Sciences and the University Research Facility in Chemical and Environmental Analysis at The Hong Kong Polytechnic University for the technical and instrumental support.

## Additional information

### Author contributions

Chengxi Liu: Conceptualization, Formal analysis, Investigation, Methodology, Validation, Visualization; Writing – original draft. Qinyu Jia: Investigation, Methodology, Validation, Visualization; Writing – review & editing. Chang Zhao: Investigation, Methodology, Validation, Writing – review & editing. Zhongping Yao: Conceptualization, Funding acquisition, Methodology, Project administration, Resources, Supervision, Writing – review & editing.

## Additional files

### Supplementary files

Supporting information includes structural alignments, mass spectra of native MS and HDX-MS, deuterium uptake lines and Woods plots, LC-MS/MS results, and MD simulations (Supplementary Figures S1-S16). Summaries of HDX experimental details (Table S1).

### Data and code availability

All data supporting this study are available within the main article and supplementary information. This study did not generate any datasets or original code.

