## Supporting Information for "SARS-CoV-2 Main Protease Inhibition Through Dimerization Promotion by Peptidomimetic Inhibitors and Disruption by Ebselen"

**Contents:**

Tables S1
Figures S1-S16

**Table S1.** Summary of the HDX experiments

| | $M^{pro}$ | | | | | | | |
|---|---|---|---|---|---|---|---|---|
| | **Free state** | **+PF-07321332** | **+PF-00835231** | **+GC376** | **+boceprevir** | **+carmofur** | **+ebselen** | **+MR6-31-2** |
| **HDX reaction details** | 10 mM phosphate buffer in 90% $D_2O$, pD 7.4, room temperature | | | | | | | |
| **Molar ratio (inhibitor/enzyme)** | - | 3:1 | | | | | | |
| **HDX time (min)** | 1, 10, 60 | | | | | | | |
| **Number of peptides** | 104 | 88 | 102 | 104 | 98 | 97 | 103 | 62 |
| **Sequence coverage** | 94.4% | 94.4% | 94.4% | 94.4% | 94.4% | 94.4% | 94.4% | 94.4% |
| **Average peptide length/redundancy** | 12.85/4.62 | 13.23/4.03 | 12.79/4.52 | 12.85/4.62 | 12.71/4.31 | 13.23/4.44 | 13.21/4.71 | 12.50/2.74 |
| **Replicates (technical)** | 3 | | | | | | | |
| **Repeatability** | 0.035 | 0.034 | 0.061 | 0.040 | 0.029 | 0.049 | 0.042 | 0.068 |
| **Significant difference in HDX (99% CI)** | - | 0.29 Da | 0.24 Da | 0.33 Da | 0.21 Da | 0.32 Da | 0.29 Da | 0.36 Da |

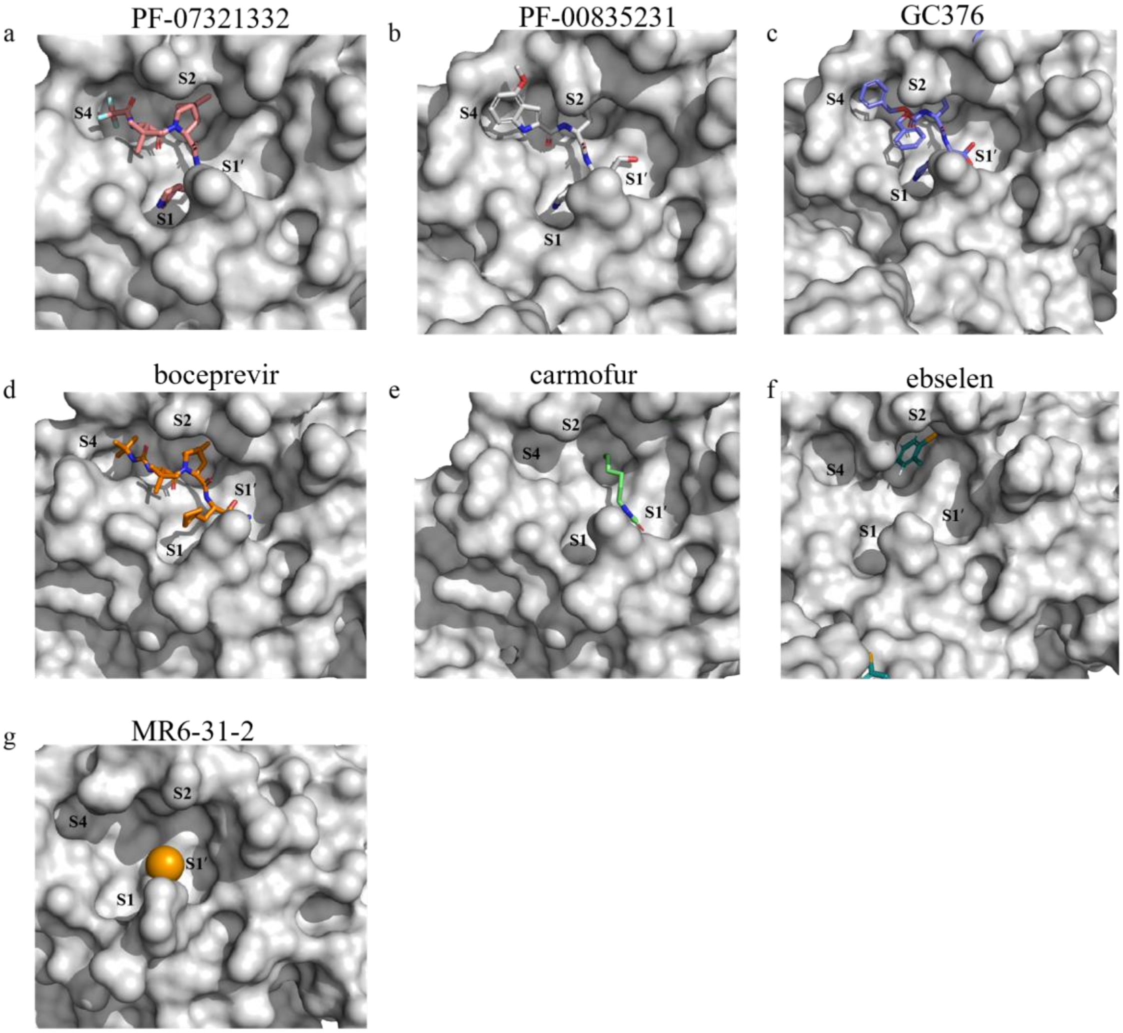


Figure S1. Surface views showing the key subpockets of SARS-CoV-2 M$^{pro}$ with: (a) PF-07321332 (PDB entry 8DZ2). (b) PF-00835231 (PDB entry 6XHM). (c) GC376 (PDB entry 7D1M). (d) boceprevir (PDB entry 7C6S). (e) carmofur (PDB entry 7BUY). (f) ebselen (PDB entry 7BFB; the orange part represents the Se atom) and (g) MR6-31-2 (PDB entry 7BAL; the orange sphere represents the Se atom). The ligand structures of ebselen and MR6-31-2 are only partially resolved in the corresponding crystal structures.

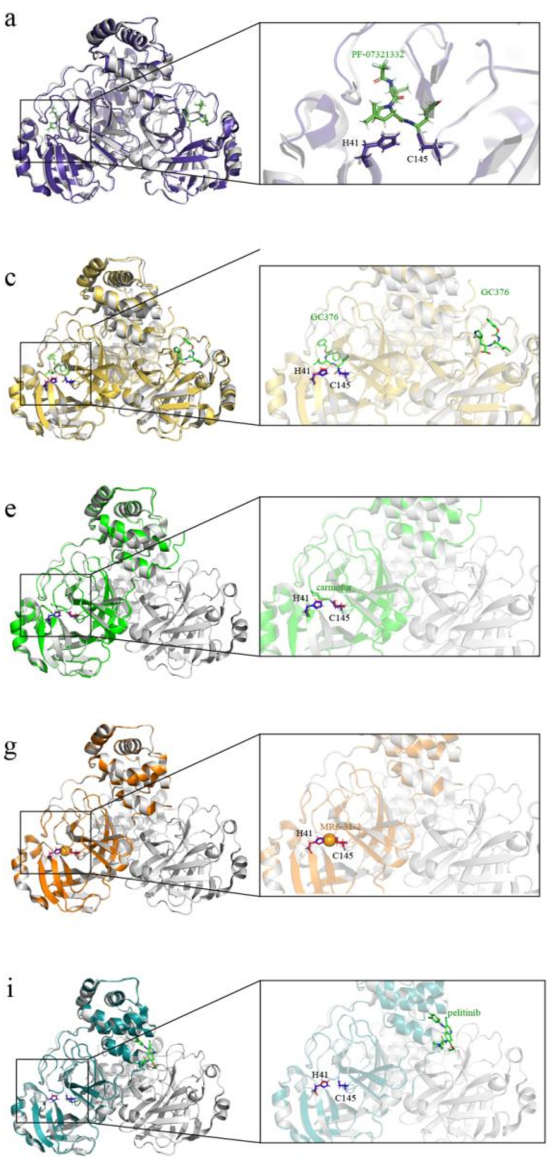


Figure S2. Alignments of the free $M^{pro}$ (PDB entry 7ALI, gray) and various inhibitor-bound $M^{pro}$ structures. (a) PF-07321332-bound $M^{pro}$ (PDB entry 8DZ2). (b) PF-00835231-bound $M^{pro}$ (PDB entry 6XHM). (c) GC376-bound $M^{pro}$ (PDB entry 7D1M). (d) boceprevir-bound $M^{pro}$ (PDB entry 7C6S). (e) carmofur-bound $M^{pro}$ (PDB entry 7BUY). (f) ebselen-bound $M^{pro}$ (PDB entry 7BFB). (g) MR6-31-2-bound $M^{pro}$ (PDB entry 7BAL, the orange sphere is the Se atom). (h) AT7915-bound $M^{pro}$ (PDB entry 7AGA) and (i) pelitinib-bound $M^{pro}$ (PDB entry 7AXM). The gray structure is an unbound state of $M^{pro}$ dimer, and the other colored structures are inhibitor-$M^{pro}$ complexes. The residues H41 and C145, and various inhibitors, are highlighted in the enlarged figures.

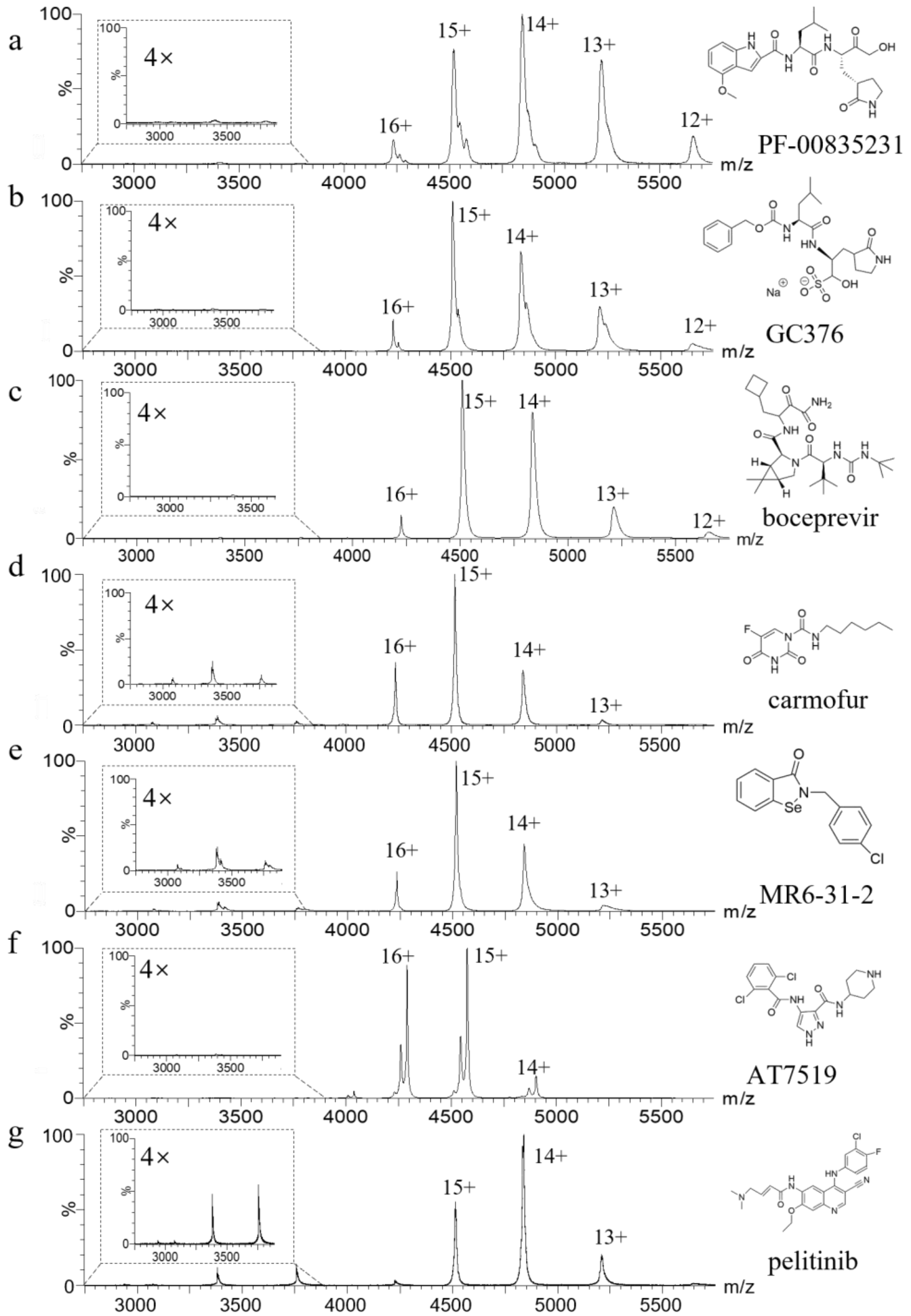


Figure S3. Native mass spectra of $M^{pro}$ (2.5 µM) in the presence of (a) PF-00835231 (7.5 µM). (b) GC376 (7.5 µM). (c) boceprevir (7.5 µM). (d) carmofur (7.5 µM). (e) MR6-31-2 (7.5 µM). (f) AT7519 (37.5 µM) and (g) pelitinib (37.5 µM).

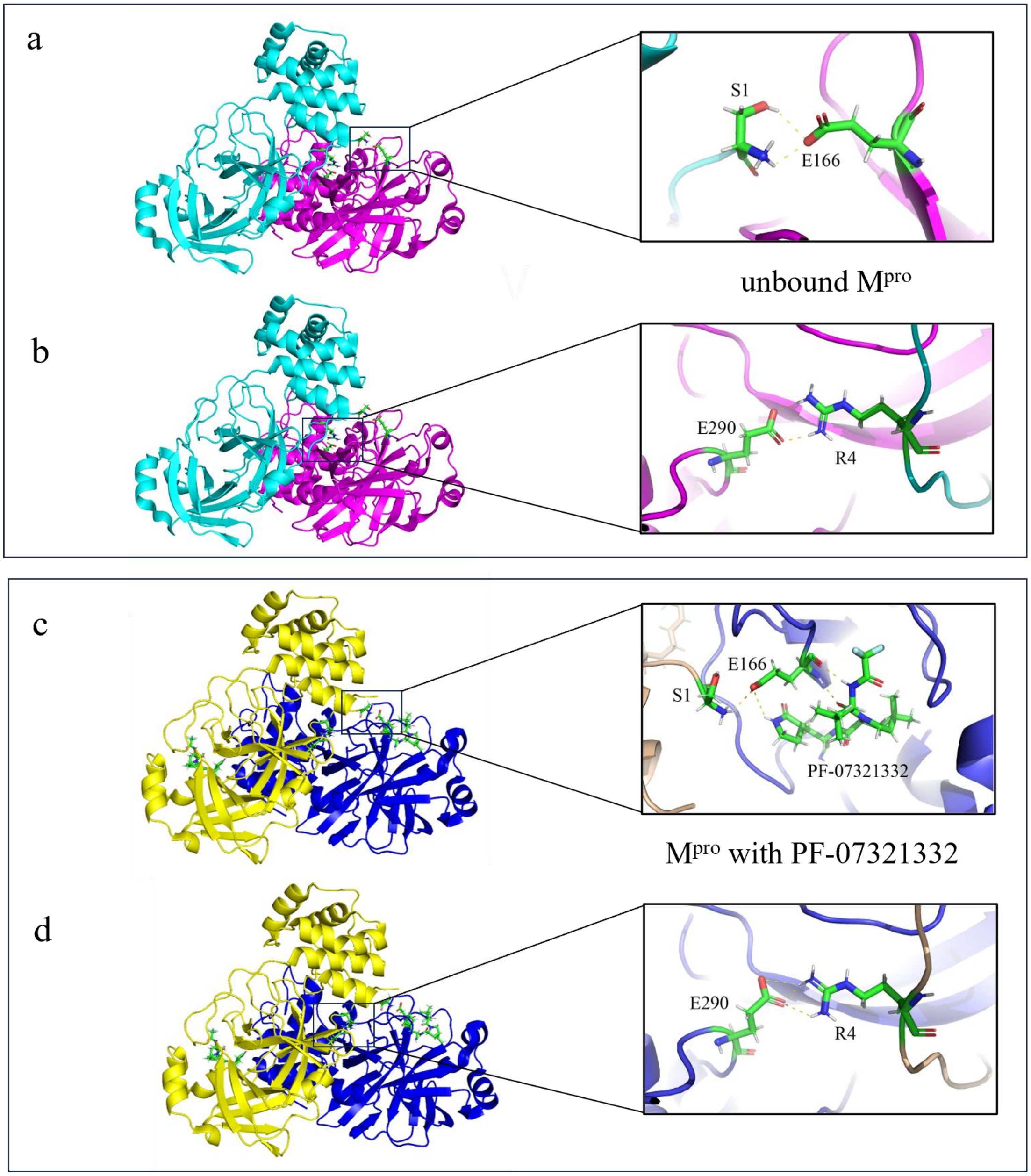


Figure S4. Hydrogen bond interactions between key residues of $M^{pro}$ dimerization. (a) Residue S1 of one monomer with E166 of the other monomer from crystallography ($M^{pro}$ alone, PDB ID 7ALI). (b) Residue R4 of one monomer with E290 of the other monomer from crystallography ($M^{pro}$ alone, PDB ID 7ALI). (c) Residue S1 of one monomer with E166 of the other monomer from crystallography ($M^{pro}$ with PF-07321332, PDB ID 8DZ2). (d) Residue R4 of one monomer with E290 of the other monomer from crystallography ($M^{pro}$ with PF-07321332, PDB ID 8DZ2).

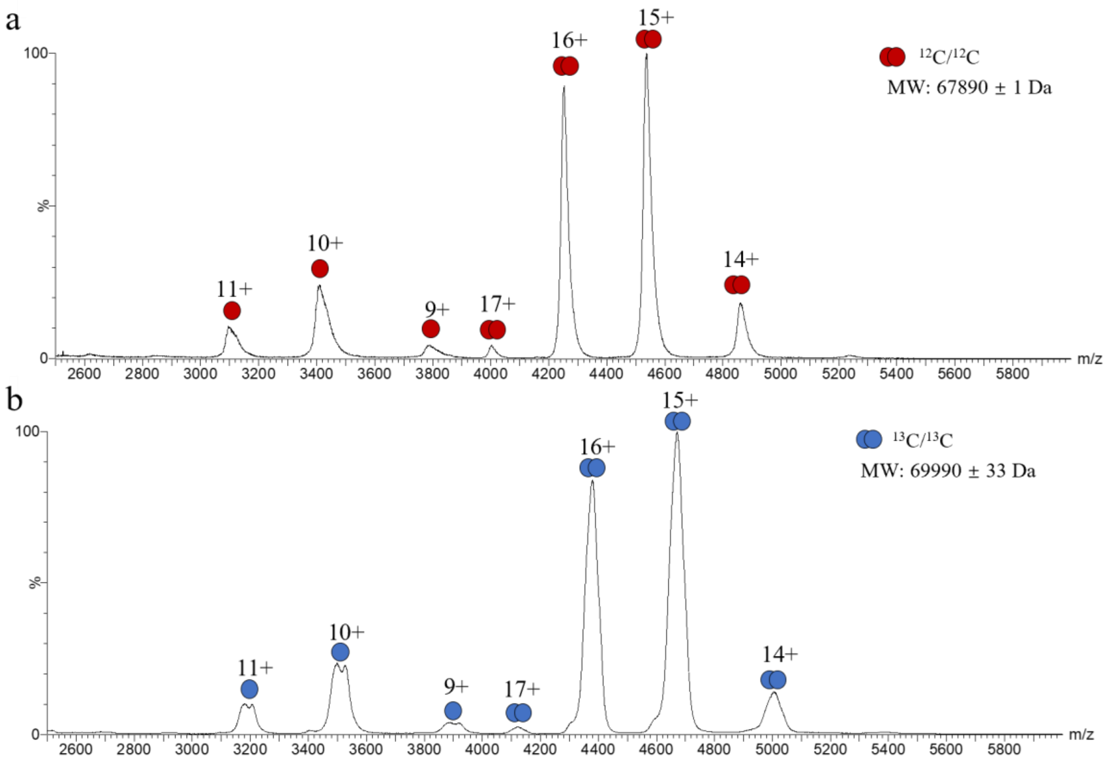


Figure S5. Characterization of $M^{pro}$ and $^{13}C$-labeled $M^{pro}$. (a) Native mass spectrum of $M^{pro}$ (2.5 μM). (b) Native mass spectrometry of $^{13}C$-labeled $M^{pro}$ (2.5 μM).

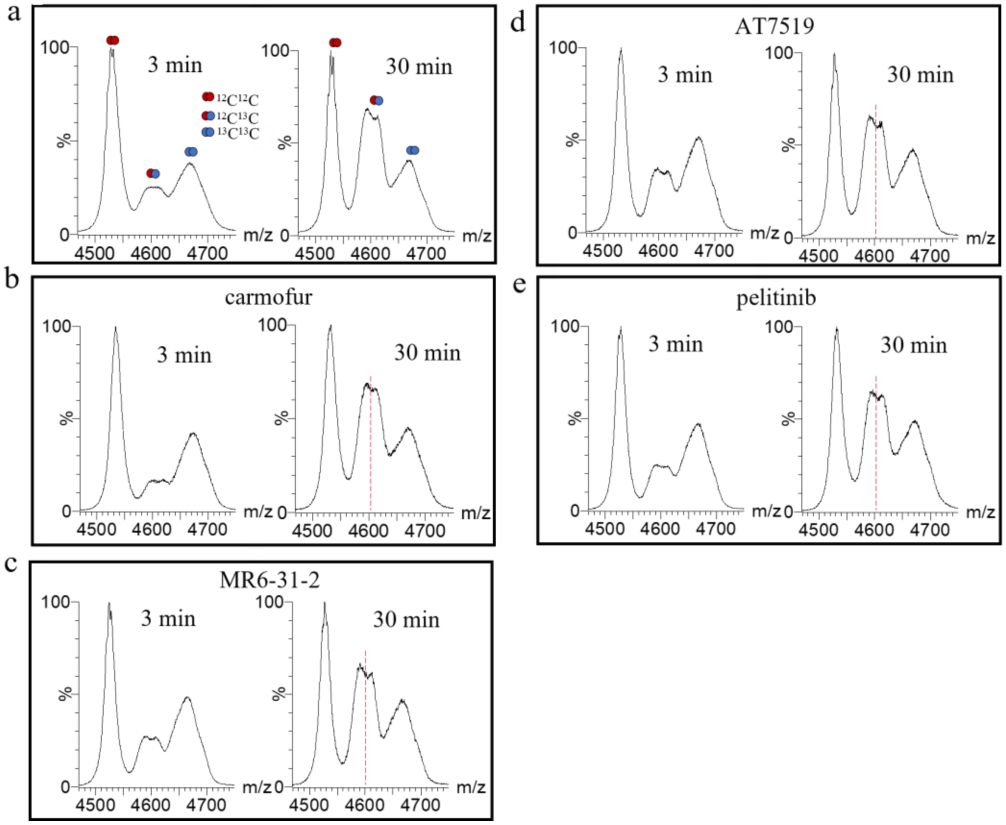


Figure S6. $M^{pro}$ subunit exchange was modulated by inhibitor-binding. Spectra were shown at the 15+ charge state of $M^{pro}$ with subunit exchange of 3 min and 30 min, respectively. (a) 2.5 µM $M^{pro}$ with 2.5 µM $^{13}$C-labeled $M^{pro}$, along with additional inhibitors: (b) carmofur (7.5 µM). (c) MR6-31-2 (7.5 µM). (d) AT7519 (7.5 µM). (e) pelitinib (7.5 µM). The red dashed lines indicate heterodimers.

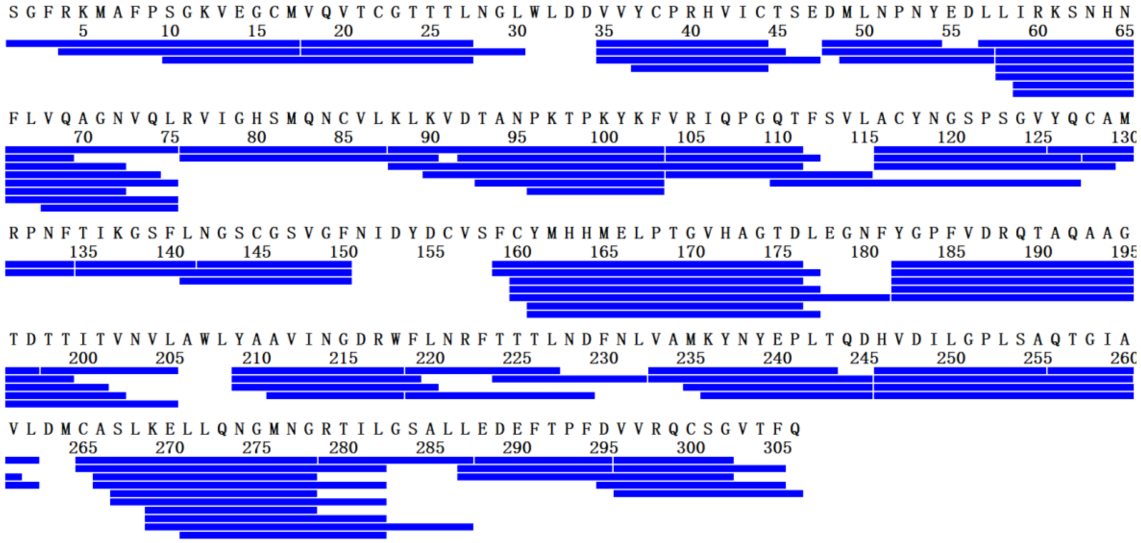


Figure S7. Coverage map of M$^{pro}$ showing 94.4% sequence coverage.

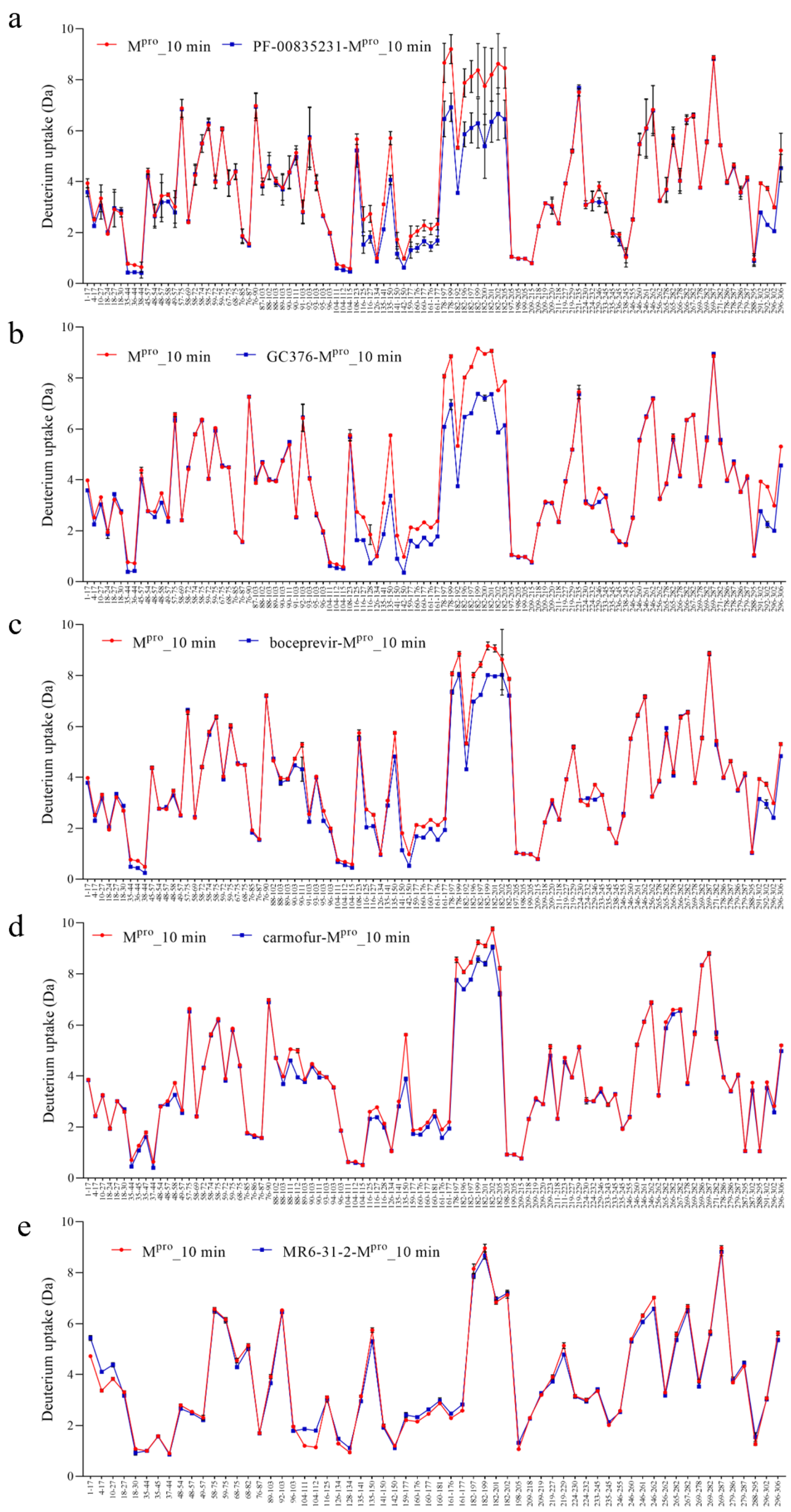


Figure S8. The deuterium uptake difference plots of $M^{pro}$ upon binding with (a) PF-00835231. (b) GC376. (c) boceprevir. (d) carmofur and (e) MR6-31-2, respectively, at 10 min.

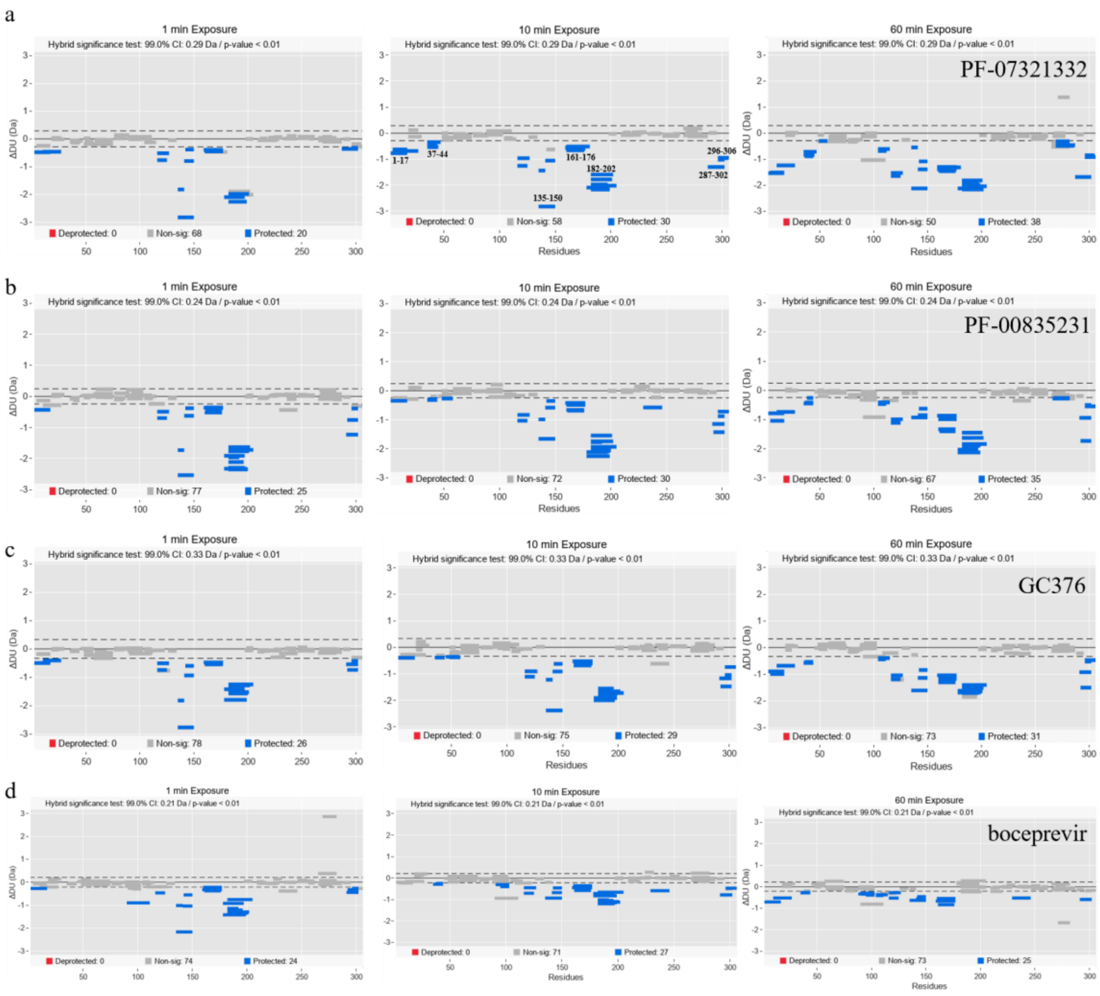


Figure S9. HDX profile of M$^{pro}$ with peptidomimetic inhibitors: (a) PF-07321332. (b) PF-00835231. (c) GC376 and (d) boceprevir. Deuterium uptake differences are presented in a Woods plot, which provides a breakdown of the peptide's ensemble for 1-, 10-, and 60-minute time points. It displays peptide length, global coverage, and deuterium uptake. A 99% confidence limit was applied to the data set to identify peptides with significant deuterium uptake. Deprotected, protected, and non-significantly different peptides are in red, blue, and grey, respectively. The presented plots have been created in Deuteros 2.0.

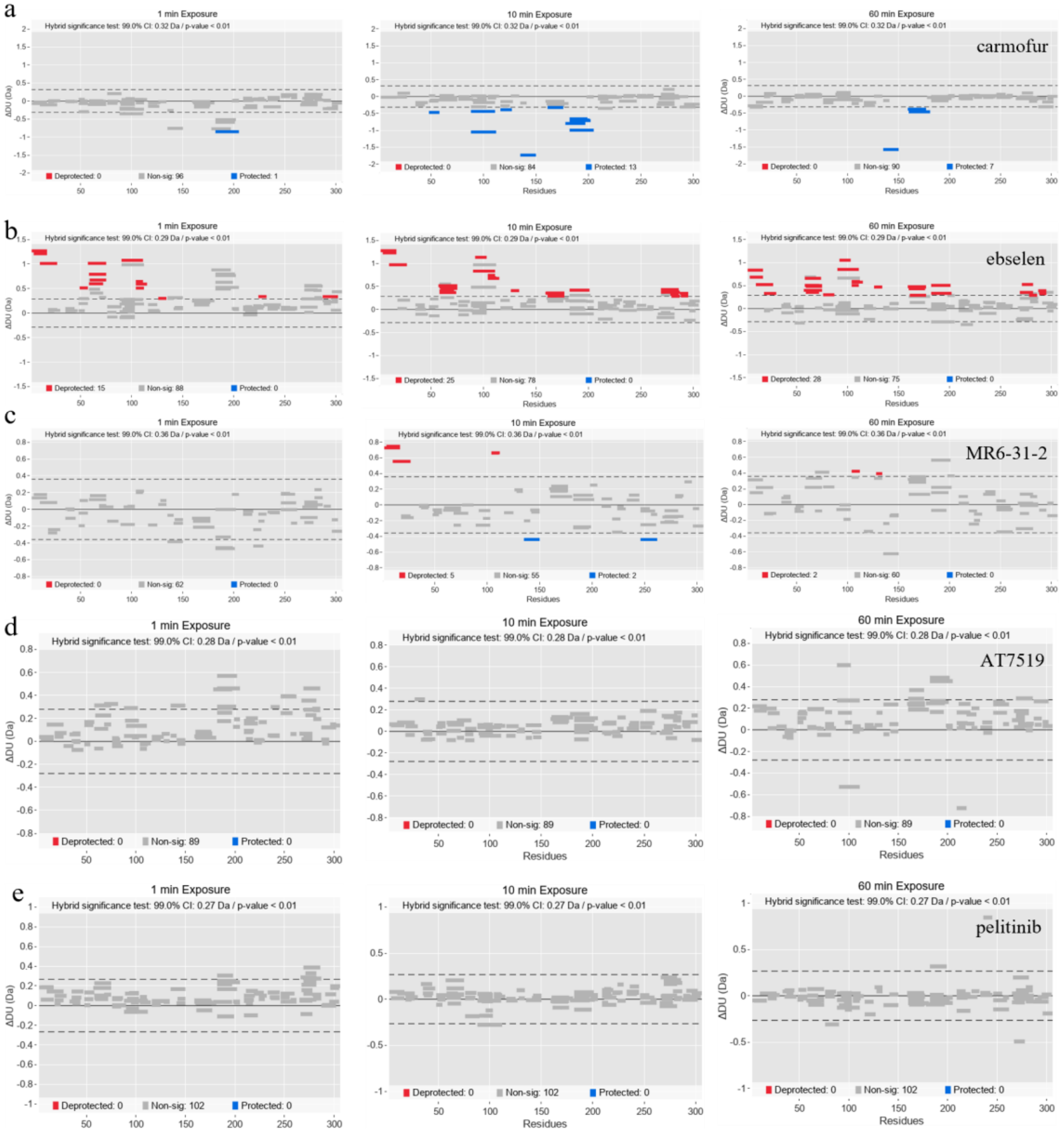


Figure S10. HDX profiles of M$^{pro}$ with inhibitors: (a) carmofur. (b) ebselen. (c) MR6-31-2. (d) AT7519 and (e) pelitinib. Deuterium uptake differences are presented in a Woods plot, which provides a breakdown of the peptide ensemble for 1-, 10-, and 60-minute time points. It displays peptide length, global coverage, and deuterium uptake. A 99% confidence limit was applied to the data set to identify peptides with significant deuterium uptake. Deprotected, protected, and non-significantly different peptides are in red, blue, and grey, respectively. The presented plots have been created in Deuteros 2.0.

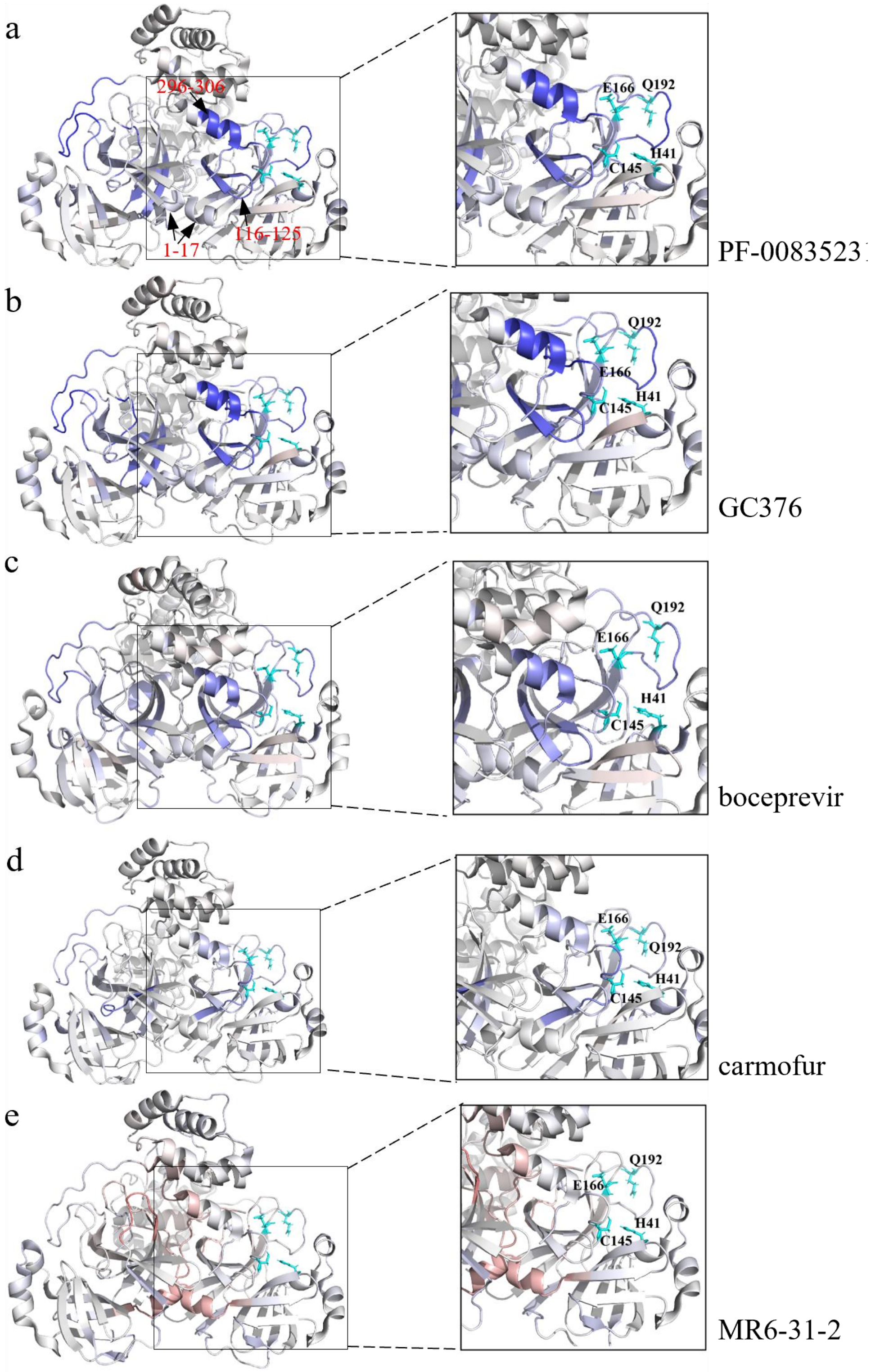


Figure S11. The deuterium uptake difference plots for all identified peptides from M$^{pro}$ upon binding with inhibitors at 10 min were labeled onto the crystal model of M$^{pro}$ (PDB ID 7ALI). (a) PF-00835231. (b) GC376. (c) boceprevir. (d) carmofur and (e) MR6-31-2.

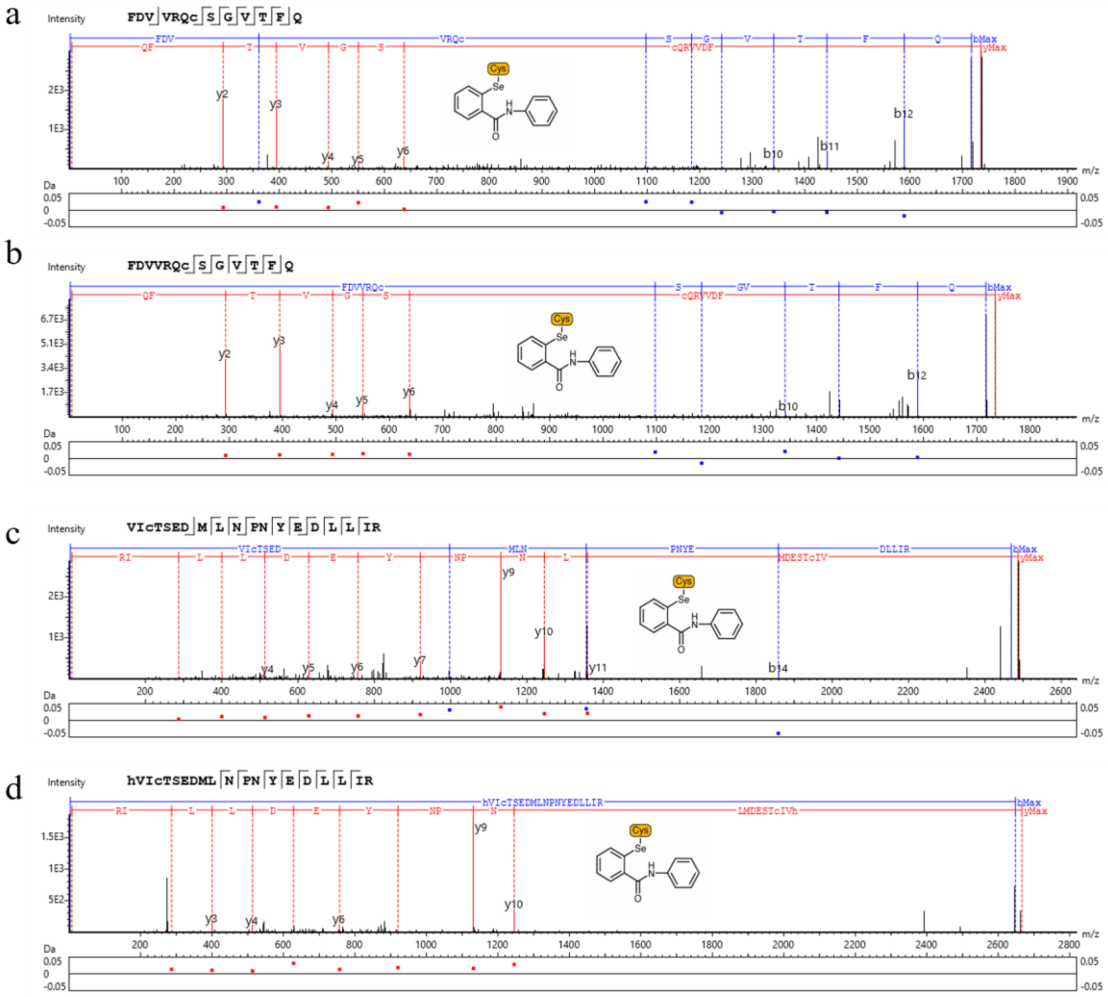


Figure S12. Tandem mass spectrometry analysis reveals that ebselen covalently bind to C44 and C300 of $M^{pro}$. (a) at *m/z* 868.8633 of the $M^{pro}$ peptide FDVVRQCSGVTFQ, containing a modification (-C13H9NOSe) induced by ebselen on C300 for $M^{pro}$ bound with ebselen at a molar ratio of 2:1. (b) at *m/z* 868.3600 of the $M^{pro}$-modified peptide FDVVRQCSGVTFQ containing a modification ($-C_{13}H_9NOSe$) induced by ebselen on C300 for $M^{pro}$ bound with ebselen at a molar ratio of 1:3. (c) at *m/z* 830.6810 of the $M^{pro}$-modified peptide VICTSEDMLNPNYEDLLIR containing a modification (-C13H9NOSe) induced by ebselen on C44 for $M^{pro}$ bound with ebselen at a molar ratio of 2:1. (d) at *m/z* 889.6963 of the $M^{pro}$-modified peptide hVICTSEDMLNPNYEDLLIR containing a modification (-C13H9NOSe) induced by ebselen on C44 for $M^{pro}$ bound with ebselen at a molar ratio of 1:3.

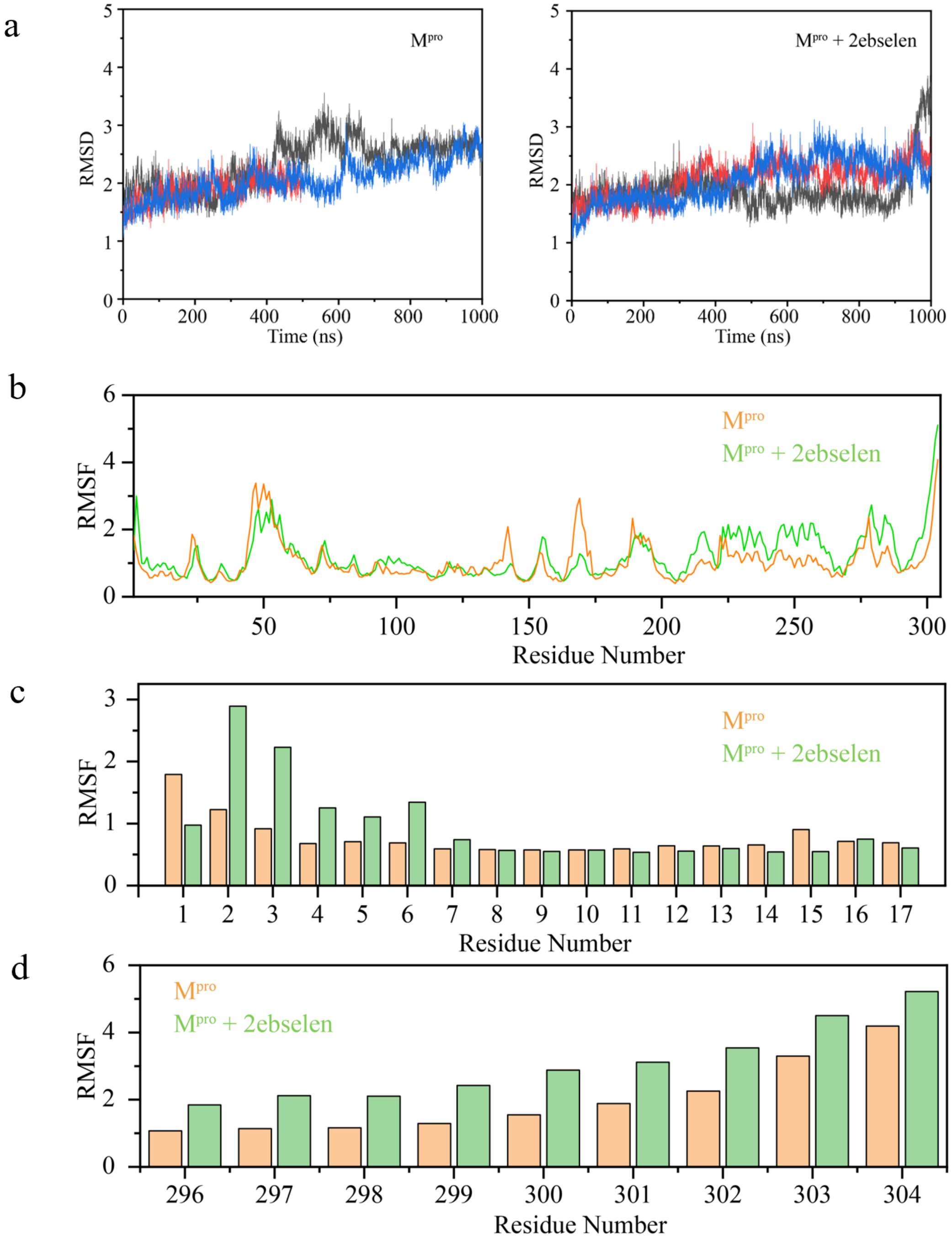


Figure S13. (a) RMSD results of unbound M$^{pro}$ and M$^{pro}$-2ebselen complex. Three colors represent the three replicates. (b) RMSF results of the unbound M$^{pro}$ protein and M$^{pro}$-ebselen complex. (c) The RMSF comparison between the N-terminal of the unbound M$^{pro}$ protein and the N-terminal of the M$^{pro}$-ebselen complex. (d) The RMSF comparison between the C-terminal of the unbound M$^{pro}$ protein and the C-terminal of the M$^{pro}$-ebselen complex.

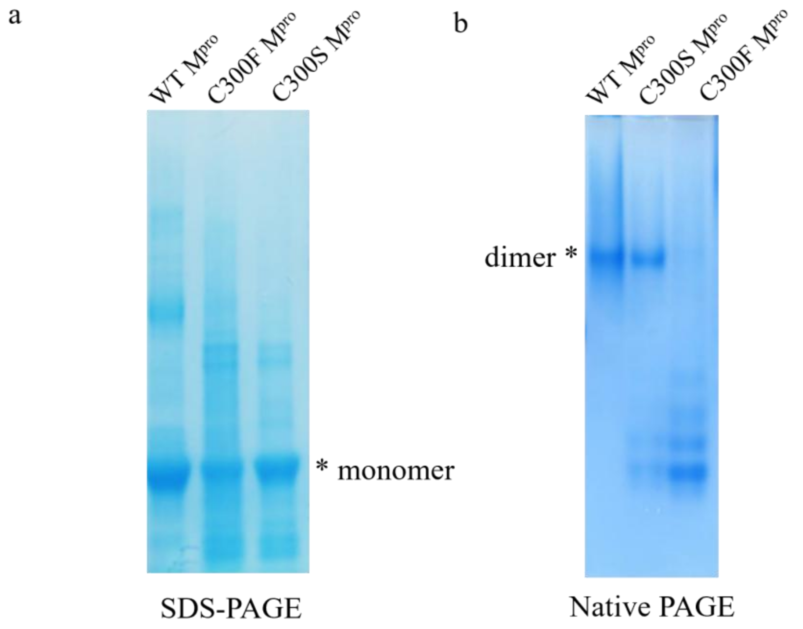


Figure S14. SDS-PAGE and native PAGE analyses of WT, C300S, and C300F M$^{pro}$. SDS-PAGE was used to assess protein integrity, whereas native PAGE was used to compare oligomeric states under non-denaturing conditions.

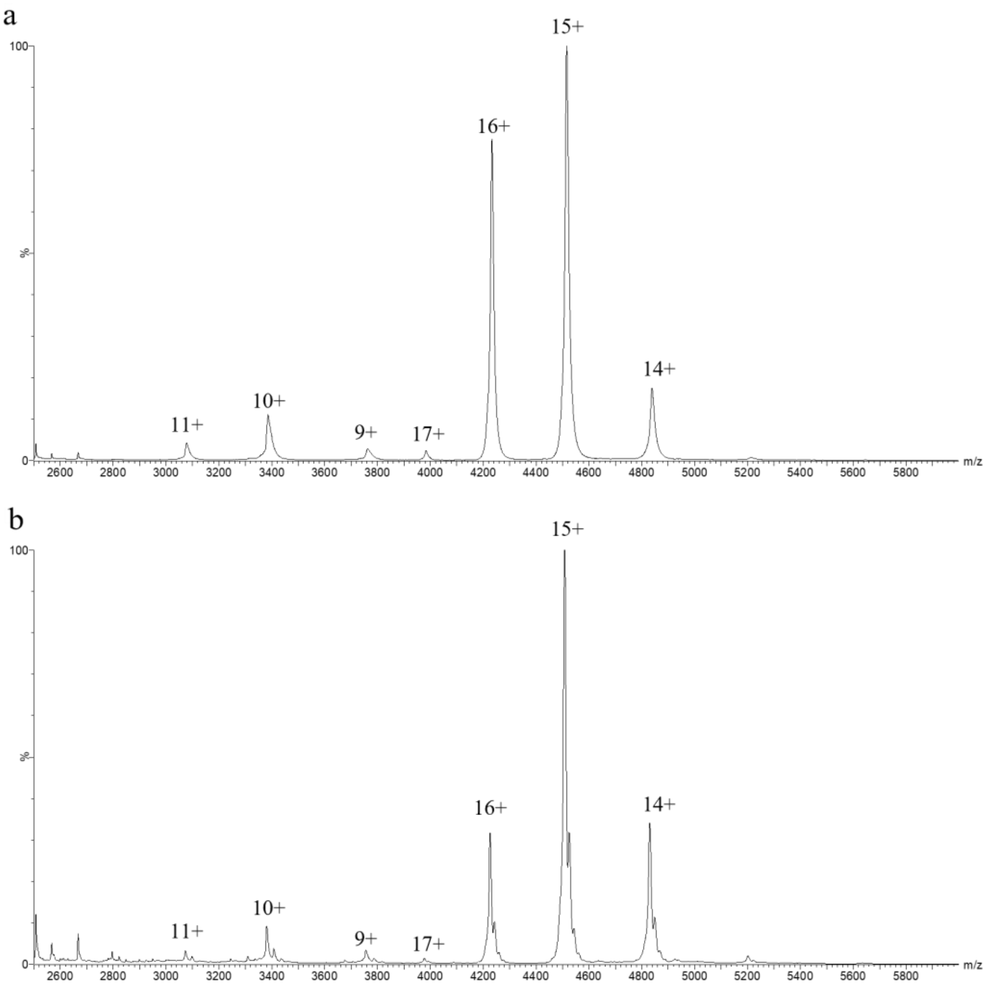


Figure S15. Native mass spectra of C300S $M^{pro}$ in the (a) absence of ebselen and (b) presence of ebselen at a 1:3 $M^{pro}$-to-ebselen molar ratio. Ebselen-bound monomeric and dimeric species were detected, whereas no marked redistribution toward the monomeric state was observed.

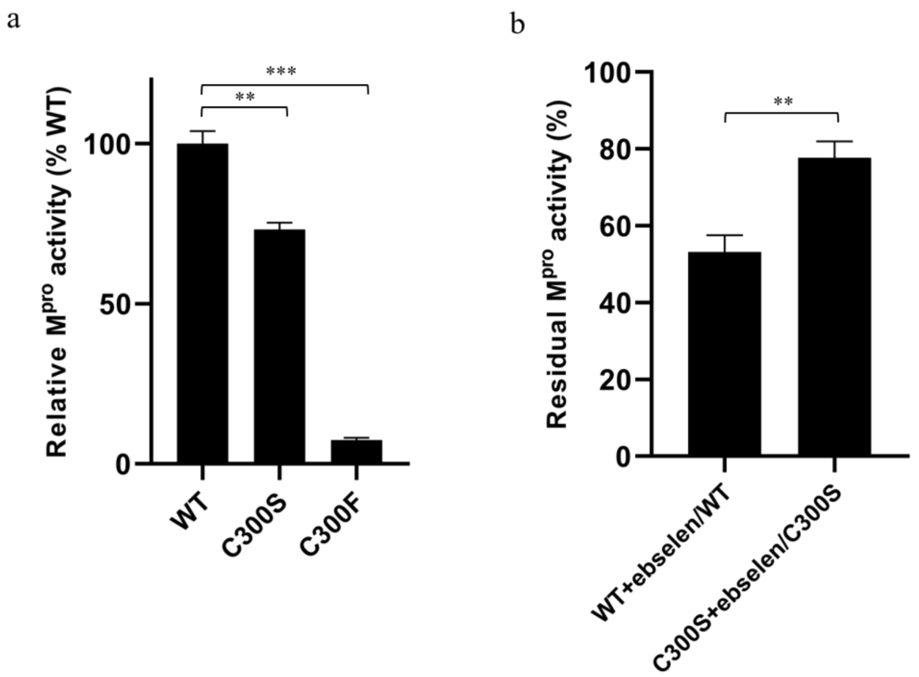


Figure S16. Effects of C300 mutations on $M^{pro}$ proteolytic activity and ebselen sensitivity. (a) Proteolytic activities of WT, C300S, and C300F $M^{pro}$ variants. Activities were normalized to WT $M^{pro}$ (100%). Data are presented as mean ± SD from three independent experiments. (b) Differential sensitivities of WT and C300S $M^{pro}$ to ebselen inhibition. Activities in the presence of ebselen were normalized to the corresponding untreated enzyme. Data are presented as mean ± SD from three independent experiments.
